# A physical theory of movement in small animals

**DOI:** 10.1101/2020.08.25.266163

**Authors:** Jane Loveless, Alastair Garner, Abdul Raouf Issa, Ruairí J. V. Roberts, Barbara Webb, Lucia L. Prieto-Godino, Tomoko Ohyama, Claudio R. Alonso

## Abstract

All animal behaviour must ultimately be governed by physical laws. As a basis for understanding the physics of behaviour in a simple system, we here develop an effective theory for the motion of the larval form of the fruitfly Drosophila melanogaster, and compare it against a quantitative analysis of the real animal’s behaviour. We first define a set of fields which quantify stretching, bending, and twisting along the larva’s antero- posterior axis, and then perform a search in the space of possible theories that could govern the long-wavelength physics of these fields, using a simplified approach inspired by the renormalisation group. Guided by symmetry considerations and stability requirements, we arrive at a unique, analytically tractable free-field theory with a minimum of free parameters. Unexpectedly, we are able to explain a wide-spectrum of features of Drosophila larval behaviour by applying equilibrium statistical mechanics: our theory closely predicts the animals’ postural modes (eigenmaggots), as well as distributions and trajectories in the postural mode space across several behaviours, including peristaltic crawling, rolling, self-righting and unbiased substrate exploration. We explain the low-dimensionality of postural dynamics via Boltzmann suppression of high frequency modes, and also propose and experimentally test, novel predictions on the relationships between different forms of body deformation and animal behaviour. We show that crawling and rolling are dominated by similar symmetry properties, leading to identical dynamics/statistics in mode space, while rolling and unbiased exploration have a common dominant timescale. Furthermore, we are able to demonstrate that self-righting behaviour occurs continuously throughout substrate exploration, owing to the decoupling of stretching, bending, and twisting at low energies. Together, our results demonstrate that relatively simple effective physics can be used to explain and predict a wide range of animal behaviours.

## Introduction

Animals have evolved a wide variety of strategies for locomotion. This allows them to search for resources, adequate habitats, or mating partners, as well as escape predators. Yet, despite this diversity, all animal movement is fundamentally a physical act, and must therefore be governed by physical laws. Determining the form of these laws, which must emerge from the interaction of nervous system, musculature, body mechanics, and the environment [1, 2], is a fundamental and inspiring problem in modern biology [3], with applications in engineering fields such as robotics [4, 5].

Recent work has leveraged the development of low-cost, high-detail imaging technologies and advanced data analysis techniques to provide fine-grained characterisations of animal behaviour. For instance, pioneering work on the nematode worm *C. elegans* demonstrated the existence of a low-dimensional postural basis in which to describe the animals’ shape changes over time [6], and similar bases have been found for other slender-bodied animals including zebrafish [7], snakes [8], and the *Drosophila melanogaster* larva [9]. Subsequent work, particularly in the nematode, has exploited such low-dimensional postural bases to theoretically investigate a wide range of behaviours using data-driven methods [6, 10–13]. However, principled, interpretable theories of animal behaviour are lacking, and it remains unclear *why* animal behaviour often appears to be low-dimensional at all, or what factors determine the dynamics and statistics within the low-dimensional postural mode space.

Here, we address these questions by studying the Drosophila larva. The larva has emerged as an excellent model system to investigate the molecular, cellular, neural, and physical bases of behaviour [14–19]. This small animal (≈ 1 mm long) hatches and grows within decaying fruit before travelling further afield in order to pupate and metamorphosise into its adult form, and so, must be capable of complex three-dimensional movement. Correspondingly, Drosophila larvae are able to perform a wide repertoire of behaviours under experimental conditions [20], including: peristaltic crawling [21], rolling [22–24], self-righting [16, 25, 26], rearing [20], hunching [23], and digging [27–30]. Recent modelling studies have exploited this system to explore the importance of biomechanics in understanding the animal’s crawling (in one spatial dimension, 1D) [19, 31–33] and substrate exploration/taxis behaviours (in two spatial dimensions, 2D) [18, 19, 34]. Additionally, the larva provides a prototypical example of the bilaterally-symmetric, slender, soft, segmented body physics that are common to many animals, and that are increasingly being exploited in the design of robotic systems [4, 5]. A physical theory of larval behaviour should thus advance general understanding of soft physics at organismal scales, and also has engineering potential.

Here, we have developed an *effective theory* [35] describing the physics of three dimensional (3D) movement in fruitfly larvae. An effective theory provides a simplified description of the key physical principles governing a system’s behaviour at a particular scale of interest, while remaining agnostic as to the causes of this behaviour at a finer scale. This approach – widely adopted in physics – has the advantage of generating concrete predictions that can be compared to measurement on a specific system (with minimal parameter fitting), while contributing to a deeper understanding of that sytem’s behaviour. We choose this approach due to its successes in statistical field theory [36] and condensed matter physics [37], where low-energy effective theories can be used to both exploit and explain the dominance and properties of long-wavelength fluctuations (i.e. the “low-dimensionality”) in the behaviour of many non-living systems [36].

Our theory describes the stretching, bending, and torsion along the fruitfly larva’s antero-posterior (AP) axis in 3D. To construct it, we employ a simple symmetry-based approach inspired by the use of the renormalisation group within statistical and quantum field theory [36, 38, 39]. We start from an arbitrary dissipative Hamiltonian field theory for a set of fields describing deformations along the larval anteroposterior axis. By restricting our attention to low energies, we arrive at a unique quadratic Hamiltonian. To capture the effects of unmodelled “high energy” dynamics, we apply an equilibrium statistical physics hypothesis to generate predictions from our low energy theory. We show that our effective theory is able to explain and predict many features across a broad range of larval behaviours, including self-righting, peristalsis, rolling, and unbiased substrate exploration. Furthermore, by performing quantitative behavioural experiments in *live* Drosophila larvae, we demonstrate that the statistics of several real larval behaviours can be surprisingly well described using our equilibrium hypothesis. We explain the low-dimensionality of behaviour by Boltzmann suppression of higher-energy, shorter-wavelength motions, with elementary low-energy, long-wavelength motions consequently dominating the animal’s behaviour. We discuss the implications of these results for behavioural control in Drosophila larvae and other animal systems.

## 1 Results

### 1.1 An effective theory of the larval midline

To advance the understanding of the physical basis of behaviour in Drosophila larvae, we constructed a theory from “first principles”. This essentially involved a “search” in the space of possible physical theories of the postural mechanics of the animal; we arrived at a unique theory by iteratively restricting our search space using symmetry and stability requirements, and through focusing attention exclusively upon the important low-energy (long-wavelength) physics by using a form of dimensional analysis inspired by the renormalisation group [36]. This type of approach has been successfully used in statistical mechanics [40], quantum field theory [38, 39], and condensed matter [37]. Perhaps most famously, Type-I superconductivity was initially formalised using the phenomenological Landau-Ginzburg theory [36], which was built using relatively simple symmetry and stability requirements, long before it was possible to investigate the microscopic properties of superconductors. It is in this spirit that we have approached larval behaviour, attempting to determine the relevant effective physics with minimal reference to microscopic details of the animal’s nervous system, musculature, or biomechanics.

We include a more detailed derivation of our theory in Appendix A, but, in essence, the approach consisted of the following steps: first, we focus our attention upon the anteroposterior (AP) axis of the animal, which for brevity we shall refer to as the *midline* (and which roughly corresponds to the neutral axis studied in rod mechanics). This choice allows us to capture the kinematics of several larval behaviours, while remaining at a tractable scale: segmental axial compression/expansion [21] and transverse bending along the midline [23, 24, 41, 42], and rotation of each body segment about the midline [16, 23–26].

Second, we introduce a reference frame with which to measure deformations of the midline (i.e. larval postures). It is necessary to choose an “undeformed” reference configuration against which to measure the deformations of the larva. For this purpose, we choose a configuration in which the larva is “straightened out” and “untwisted”. This allows us to parametrise deformations by a single spatial parameter, the arc-length *s* measured along the midline from tail (*s* = 0) to head (*s* = 1) in the reference configuration, along with the time *t* measured relative to an arbitrary initial time *t*_0_; we will often refer to *s* simply as “space”, since together with *t* it forms the underlying 2-dimensional space-time of our theory. We introduce a right-handed Cartesian frame aligned with the anatomical frame of the reference configuration. In particular, the origin is fixed at the tail; the positive *x*-axis is directed along the midline and passes through the centroid of each transverse cross-section of the body, at the intersection of the anatomical sagittal and coronal planes; the *x — z* plane is aligned with the sagittal plane of the reference configuration with the positive *z — axis* along the dorsal axis; the *x — y* plane is aligned with the coronal plane with the positive *y — axis* directed towards the left-hand-side of the larva. The deformation of the larva is then given by the fields *x*(*s, t*), *y*(*s, t*), *z*(*s, t*) which measure the time-varying Cartesian displacement at each point along the midline, and *θ*(*s, t*) which measures the rotation of transverse cross sections about the midline, relative to the undeformed configuration (Figure 1A). For brevity we combine these four scalar fields into a vector *ψ* = [*x, y, z, θ*]^*T*^ (*s, t*).

**Fig 1.**
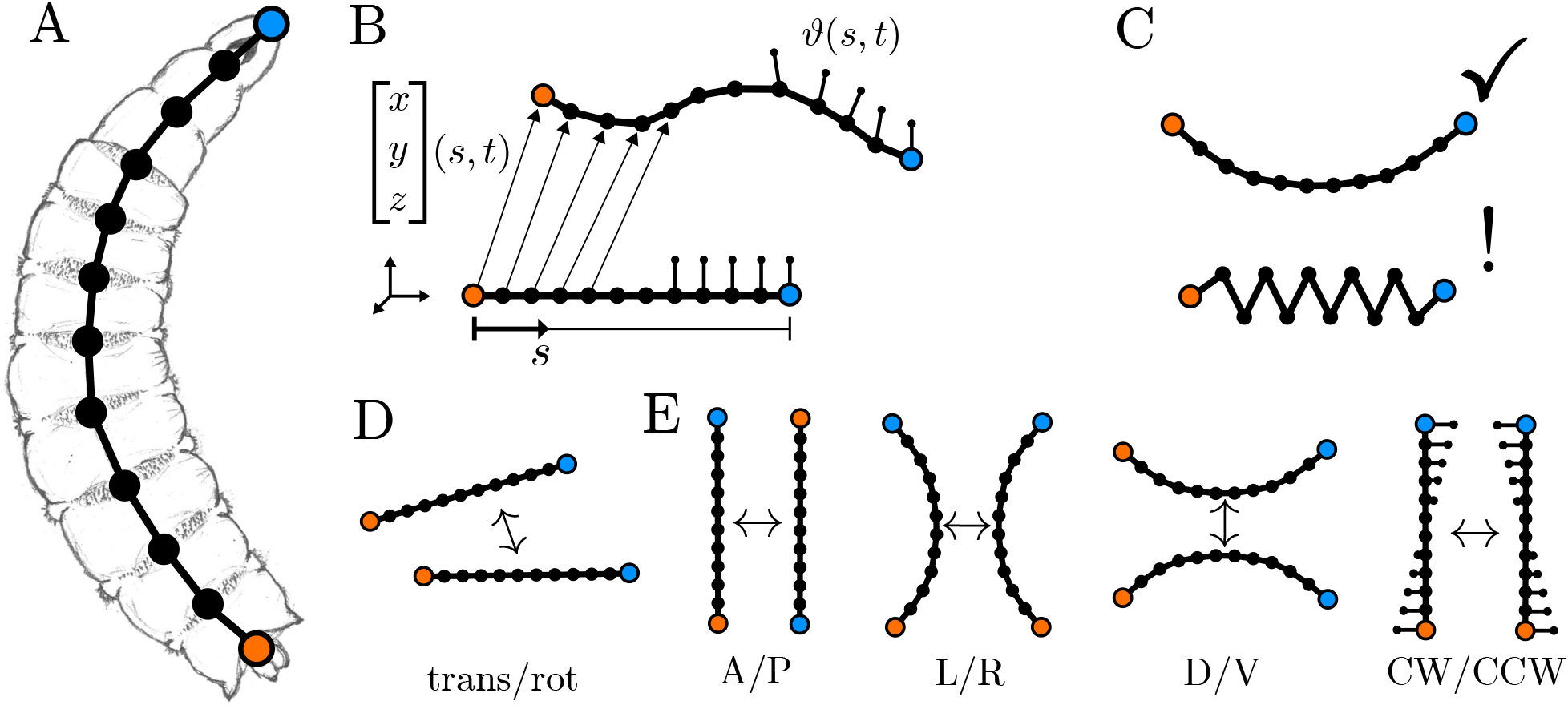
Constructing an effective theory of larval midline physics. A) An illustration of the *Drosophila* larva with its midline overlaid (black, blue=head, orange=tail). B) We measure the displacement of the larva from an undeformed configuration (bottom) to a deformed configuration (top) via four scalar fields parametrised by arc-length *s* (measured along the body from *s* = 0 at the tail, to *s* = 1 at the head) and time *t*. The fields correspond to the 3 translations *x*(*s, t*), *y*(*s, t*), *z*(*s, t*) and the torsional rotation of the body about each point *θ*(*s, t*). C) We restrict attention to low energy, long wavelength, low frequency motions of the midline (top) rather than the higher energy, short wavelength, high frequency motions (bottom). D) We impose continuous symmetry of the effective theory under overall translations and rotations. E) We impose discrete symmetry of the effective theory under anterior–posterior (A/P) axial reflections, left-right (L/R) mediolateral reflections, dorsal–ventral (D/V) reflections, and clockwise–counterclockwise (CW/CCW) torsional reflections. We also require that our effective theory be described by an analytic, local Hamiltonian field theory (not illustrated in this figure).

Thirdly, we use of an effective statistical equilibrium hypothesis, postulating that the probability of a given field configuration *ψ* can be expressed in terms of an effective postural mechanical energy *H* [*ψ*] (also referred to as the Hamiltonian) via the Boltzmann distribution

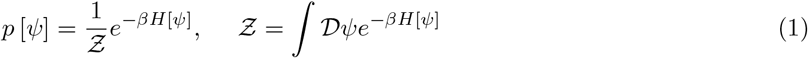

Here *Ƶ* is the partition function, which is defined so as to normalise the probability distribution. Because we are dealing with continuous fields the partition function takes the form of a path integral, with the measure 𝒟*ψ* intended to denote integration over all possible field configurations. We assume that the Hamiltonian *H* has the locality property, meaning that it can be written as an integral over space *s* of a Hamiltonian density *ℋ* depending on the field *ψ* and its derivatives with respect to *s* and *t* as 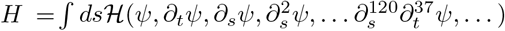. For brevity we will often refer to the Hamiltonian density as simply the Hamiltonian; although this terminology is technically incorrect it is in common usage within field theory [36, 38, 39].

As in many other statistical field theories [36], the effective form of the Hamiltonian density *ℋ* is strongly constrained by symmetry properties, and takes a very simple form in the limit of low energies. In particular, since the larva moves at speeds which are almost negligible relative to the speed of light, we use a non-relativistic classical mechanics framework and impose Galilean invariance. This means that the energy of a given larval postural configuration should not depend on overall translations and rotations of the larva in space. Mathematically, this manifests as a minimum order of differentiation required for a term to be present in the Hamiltonian, e.g. the Hamiltonian cannot contain a term in *x*^2^ because on translation *x* → *x* +Δ*x* we have *x*^2^ → (*x* +Δ*x*)^2^ ≠ *x*^2^, however the Hamiltonian can contain a term in (*∂*_*s*_*x*)^2^ because *∂*_*s*_(*x* +Δ*x*) = *∂*_*s*_*x*.

We further impose reflection symmetry; this is motivated by the bilateral symmetry of the larva, which we interpret as meaning that a left-handed deformation of the midline should have the same energy as a right-handed deformation (so that the transformation *y* → *−y* should leave the Hamiltonian invariant). Combining this left-right symmetry with overall rotational symmetry generates a greater range of reflection symmetries – if we rotate the larva 180° in the sagittal plane before performing a left-right reflection, we have, in effect, performed a dorsal/ventral reflection, and we can clearly see that the Hamiltonian should be invariant under such D/V reflections *z* → *−z*; similar constructions lead to requirements for invariance under axial reflections *x* → *−x* and clockwise/counterclockwise reflections *θ* → *−θ*. Incorporating these reflection invariances into the Hamiltonian density decouples the scalar fields *x, y, z, θ* from one another up to quadratic order, and further removes all linear terms from the Hamiltonian, so that we can write it in the form

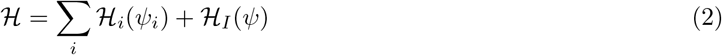

where dependence on the derivatives of fields are implied, the term *ℋ*_*I*_ contains all quartic and higher order terms, and the summation is over a set of purely quadratic, decoupled Hamiltonians, one for each field *x, y, z, θ* ∈ *ψ*.

As we restrict attention to low energies (long wavelengths, low frequencies, and small field/derivative amplitudes) both the higher order terms in *ℋ*_*I*_ and the higher derivatives within the quadratic Hamiltonians *ℋ*_*i*_ become less relevant, and the Hamiltonian is dominated by the lowest order derivatives allowed by our symmetry requirements (Appendix A). In this case we therefore arrive at Hamiltonian densities of fixed form

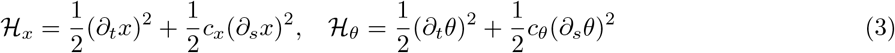

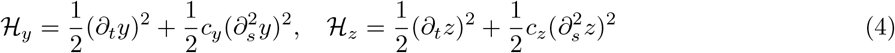

where we have eliminated the coefficients of the first term in each Hamiltonian via a suitable rescaling of the fields, in order to leave just four free parameters *c*_*i*_. We note in passing that these Hamiltonians are in the same form as the Hamiltonian densities for stretching, bending, and torsion of a deformable rod in classical linear elasticity theory, with stretching governed by the linear wave equation and bending governed by the Euler-Bernoulli beam theory [43]. Our theory is thus strongly reminiscent of the phenomenological elasticity models that have been used with great success in understanding mechanical properties of DNA and other polymers [40, 44–46]; the key difference is our inclusion of kinetic energy, owing to the larva’s relatively large size and our interest in the larva’s dynamics.

Having decoupled the Hamiltonian, the Boltzmann distribution and the partition function now factor into separate contributions from each field

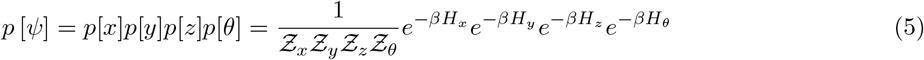

with 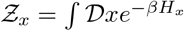, and equivalent expressions for the other field-specific partition functions.

Since, by necessity, behavioural data tracks only discrete points along the larval body, we next discretise the scalar fields and their derivatives using standard finite difference schemes. We choose to measure the fields at *N* = 12 discrete points along the midline, corresponding to the boundaries between the 11 body segments in the Drosophila larva: T1 – T3 (thoracic), A1 – A8 (abdominal) along with the head and tail extremities. It is necessary to introduce boundary conditions on our scalar fields in order to correctly specify the particular difference operators used in our discretisation. We impose vanishing force/moment (Neumann) boundary conditions on the transverse and torsional fields, and periodic boundary conditions on the axial field. The Neumann boundary conditions represent the absence of bending or twisting moments at the head and tail extremities, while the periodic axial boundary conditions represent the presence of a *visceral piston* in the larva which couples the motion of the head and tail.

We reduce the number of free parameters in our model by assuming that the transverse fields have an “internal” circular rotational symmetry, i.e. we assume that the physics of mediolateral and dorsoventral motions should be equivalent and write *c*_*y*_ = *c*_*z*_, leaving three free parameters. This is motivated both by a desire for simplicity in our theory, and by the approximately circular cross-section of the larva, which should lead to a similar passive bending response in the mediolateral and dorsoventral directions. The introduction of periodic boundary conditions to model the visceral piston of the larva introduces a similar “rotational” symmetry into the effective theory. We discuss the implications of these symmetries later in the paper, during our modal analysis and while studying peristalsis and rolling behaviours.

Our discretised Hamiltonian is then given by

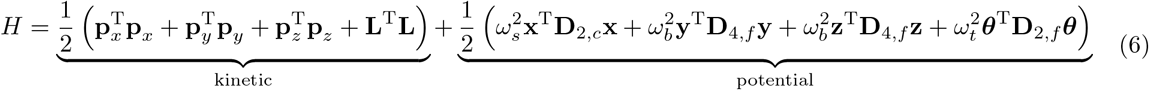

where we have once again combined the Hamiltonians for the separate fields, i.e. *H* = *H*_*x*_ + *H*_*y*_ + *H*_*z*_ + *H*_*θ*_, and we have indicated that the Hamiltonian corresponds to the total effective mechanical energy of the midline, and is the sum of kinetic and potential energies.

In these expressions the lower-case bold symbols **x, y, z, *θ*** denote vectors containing measurements of the scalar fields *x, y, z, θ* at uniformly sampled discrete points along the midline *s*_*i*_, while the vectors **p**_*x*_, **p**_*y*_, **p**_*z*_ contain the discrete measurements of the translational momenta and **L** contains discrete measurements of angular momentum about the midline.

The parameters *ω*_*s*_, *ω*_*b*_, *ω*_*t*_ measure the ratio of elastic to inertial effective forces and determine the absolute frequencies of stretching, bending, and twisting motions, respectively. Note these parameters are not numerically equal to the parameters *c*_*i*_ in the earlier Hamiltonian density, as they have absorbed constants associated with the discretisation of space. The upper-case bold symbols **D**_2,*c*_, **D**_4,*f*_, **D**_4,*f*_, and **D**_2,*f*_, denote the finite difference operators we mentioned above. **D**_2,*c*_ is the (*N −* 1) × (*N −* 1) second difference matrix with periodic boundary conditions

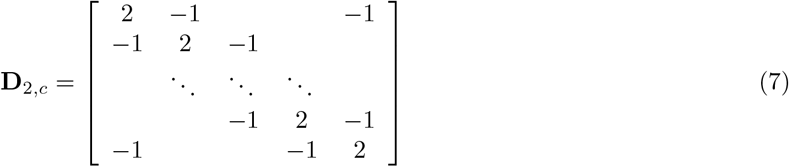

**D**_2,*f*_ is the *N* × *N* second difference matrix with free-free boundary conditions

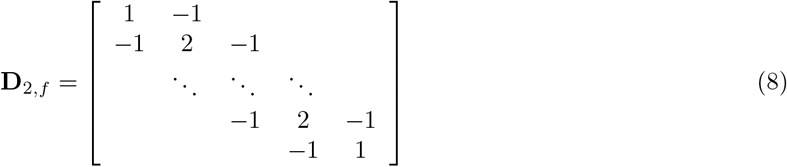

and **D**_4_ is the *N* × *N* fourth difference matrix with free-free boundary conditions

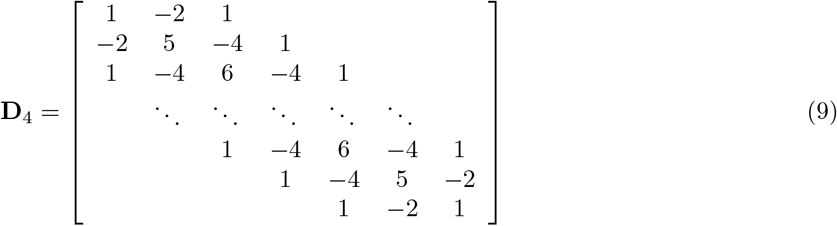

### 1.2 Neuromuscular and dissipative effects

Before exploring the behaviour of our effective theory, we first consider the effects of neuromuscular forcing within the low-energy limit that we set above, since it is not intuitively clear whether our Hamiltonian approach can effectively model these effects. To examine this issue we postulate that the Hamiltonian derived above correctly encapsulates the low-energy dynamics, as well as the statistics, of our deformation fields. In this case, we can derive partial differential equations governing evolution of the fields *x, y, z, θ* ∂ *ψ* by first taking the Legendre transform of the Hamiltonian to find the Lagrangian density *∂*(*ψ*) = ∑_*i*_ *π*_*i*_*∂*_*t*_*ψ*_*i*_ *− ℋ* (*ψ*) and then finding the corresponding Euler-Lagrange equations which extremise the action *S* = ∫ *dtdsℒ*. Since our Hamiltonian density is quadratic, so is the corresponding Lagrangian density, and the Euler-Lagrange equation is therefore linear, and can be written with generality as

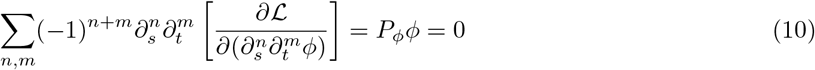

Here, *ϕ* ∂ *ψ* is used as a stand-in for any of the scalar fields *x, y, z, θ*, and *P*_*ϕ*_ is a linear differential operator encoding the Euler-Lagrange dynamics. In the presence of generalised forces acting on the field *ϕ*, the right hand side is nonzero, and in general we will have instead

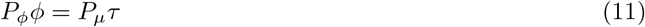

where *τ* = *τ* (*s, t*) is a field whose derivatives produce generalised forces conjugate to the mechanical field *ϕ*. Note that if our field *τ* is to produce only forces *internal* to the mechanical system, such as the forces produced by muscles which have attachment sites within the body, then, by definition, the resulting dynamics must be Galilean invariant [47] (this is equivalent to requiring that muscle forces alone cannot produce a change in the total momentum or angular momentum of the midline in the absence of a ground reaction). Just as Galilean invariance limited the lowest order derivatives that could appear in our Hamiltonian density (and by extension *P ϕ*), here this invariance limits the lowest order derivative that can appear in *P*_*µ*_. As an example, if *ϕ* represents the mediolateral transverse deflection field *y*(*s, t*) then the field *τ* (*s, t*) could represent an internal bending torque field, in which case we would have 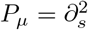. Conversely, if *ϕ* represents the axial field *x*(*s, t*) then the field *τ* (*s, t*) could represent internal tension forces, and we would have *P*_*µ*_ = *∂*_*s*_.

The field *τ* may also have its own dynamics and may be reciprocally coupled to the mechanical field *ϕ*. Assuming we remain within a linear, low-energy regime, we can write

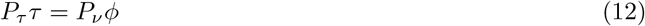

where *P*_*ν*_ is constrained by symmetry. For instance, Galilean invariance will again set the lowest order of derivative present within *P*_*ν*_, representing the idea that the neuromuscular system cannot produce forces which depend upon the overall translation or rotation of the larva in space. However, in general *P*_*τ*_, representing the dynamics of the *τ* field itself, is not required to satisfy any symmetries.

Now we note that we can apply the operator *P*_*µ*_ to Equation 12, and assuming the commutativity of our linear partial differential operators (i.e. assuming Clairaut’s Theorem holds) we can write

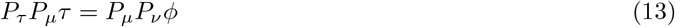

but by Equation 11 we have *P*_*µ*_*τ* = *P*_*ϕ*_*ϕ*, so this is equivalent to

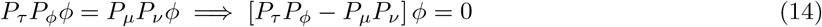

This means the field *τ* can be eliminated by modifying the dynamics for the field *ϕ*. This can be rewritten simply as

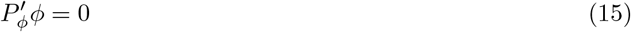

where 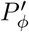 is again a linear differential operator. Note that 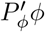 is guaranteed to be the Euler-Lagrange equation of a *new* quadratic Lagrangian *ℒ′*, although this Lagrangian may contain fractional derivatives (corresponding to odd-order derivatives in the Euler-Lagrange equations [48], see Appendix C). Crucially, the lowest order terms appearing in *ℒ*′ will generally be of the same order as the lowest order terms in *ℒ* since both *P*_*ϕ*_ and *P′*_*ϕ*_ are Galilean invariant. Thus, on rescaling, *ℒ*′ should be equivalent to *ℒ* up to a change in value of the free parameters, and thus the new Hamiltonian *ℋ′* will be similarly equivalent to *ℋ*.

We therefore conclude that, within a linear approximation to the dynamics, and upon applying a rescaling of the fields, space, and time, the only effect of neuromuscular forcing is a rescaling of the free parameters in our original Hamiltonian density. This is an essentially irrelevant difference, since the values of the free parameters are to be fixed by experiment.

### 1.3 Modal analysis and Gaussian statistics

Having formulated our effective theory, we next seek a convenient coordinate system in which to explore its behaviour. For this, we apply *modal analysis*, which, for our purposes, is formally identical to principal components analysis (PCA) (see below). Modal analysis gives a description of our theory in terms of a set of non-interacting, elementary collective motions of the body segments, with each collective motion measured by a *modal coordinate*, analogous to a principal component, and possessing a characteristic frequency and *mode shape* describing the spatial pattern of movement, equivalent to a principal component vector – also referred to as an *eigenmaggot* in the field [9]

The mode shapes are formally given by the eigenvectors of the difference matrices in our effective theory, so that transforming to modal coordinates diagonalises these matrices. This means that the Hamiltonian can be written in the form

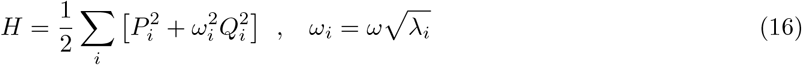

Here *Q*_*i*_ denotes the *i*’th modal coordinate, *P*_*i*_ denotes its conjugate momentum, and *ω*_*i*_ is the characteristic frequency of the mode. Frequency is written in terms of an absolute frequency scale *ω*, and the *i*’th eigenvalue *λ*_*i*_ of the relevant difference operator. The absolute frequency scale *ω* corresponds to the relevant free parameter *ω*_*s*_, *ω*_*b*_, or *ω*_*t*_ in our discretised Hamiltonian, depending upon which field the mode belongs to. Clearly the ratio of frequencies for two modes of the same deformation field is independent of the absolute frequency scale, depending only upon the eigenvalues of the difference matrices

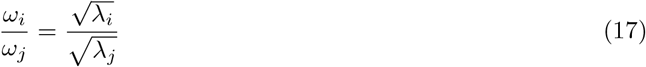

In terms of the modal coordinates, the Boltzmann distribution completely factors as

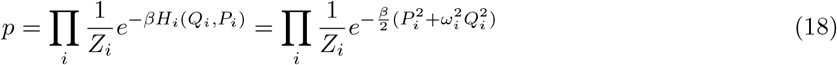

where *Z*_*i*_ = 2*π/βω*_*i*_ is the partition function for the mode described by *Q*_*i*_, *P*_*i*_. Clearly, in this description, each modal coordinate has simple Gaussian statistics. Indeed, we can interpret our modal analysis as finding the eigenvectors of the covariance matrix of a multidimensional Gaussian distribution, obtained from the Boltzmann distribution with our specific quadratic Hamiltonian. Here is the connection to PCA we described earlier – the eigenvectors of the covariance matrix are exactly the principal component vectors, which coincide, exactly, with our mode shapes; the principal components, representing projection onto these vectors, are then exactly equivalent to the modal coordinates. Because the mode shapes depend only upon the difference matrices in the Hamiltonian, we can make parameter-free predictions of the principal component vectors extracted from larval behaviour.

Furthermore, examination of Equation 18 shows that each modal coordinate (principal component) *Q*_*i*_ has variance 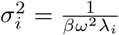, as predicted by equipartition of energy amongst the modes [40]. It is thus clear that the *proportion* of variance explained by each mode of a given deformation field is independent of the temperature *β* and absolute frequency *ω* parameters, since we have

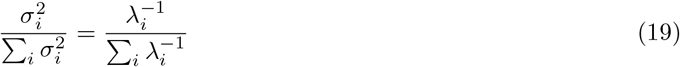

This, therefore, enables us to make parameter-free predictions of both the mode shapes and the proportion of variance explained by each mode (Table 1 and Figure 2; see Appendices E, F for calculations of the eigenvalues and eigenvectors for axial and torsional deformations, and G for analytical approximations for transverse deformations).

**Table 1.** Cumulative percentage of variance explained by the stretching, bending, and twisting modes during different larval behaviours, compared to model predictions (fw crawl = forward peristaltic crawling; bk crawl = backward peristaltic crawling; explore = unbiased substrate exploration; roll = rolling; SR = self-righting). Reported values are median across trials/individuals.

| mode | fw crawl | bk crawl | model | explore | roll | SR | model | SR | model |
| --- | --- | --- | --- | --- | --- | --- | --- | --- | --- |
| 1 | 30.70 | 27.40 | 31.50 | 85.72 | 82.01 | 90.22 | 82.67 | 41.43 | 41.14 |
| 2 | 57.91 | 53.90 | 62.99 | 94.52 | 91.60 | 96.75 | 93.91 | 59.40 | 60.06 |
| 3 | 65.43 | 63.83 | 71.55 | 97.02 | 95.84 | 98.51 | 97.03 | 70.19 | 71.23 |
| 4 | 79.34 | 78.70 | 80.10 | 98.27 | 97.52 | 98.95 | 98.28 | 77.00 | 78.84 |
| 5 | 82.81 | 83.74 | 84.48 | 98.83 | 98.36 | 99.33 | 98.93 | 83.71 | 84.56 |
| 6 | 88.98 | 89.70 | 88.85 | 99.43 | 99.15 | 99.60 | 99.32 | 88.21 | 89.17 |
| 7 | 91.92 | 93.07 | 91.88 | 99.72 | 99.60 | 99.81 | 99.60 | 90.92 | 93.12 |
| 8 | 94.17 | 94.56 | 94.90 | 99.93 | 99.91 | 99.96 | 99.81 | 95.00 | 96.67 |
| 9 | 97.38 | 97.56 | 97.45 | 100.00 | 100.00 | 100.00 | 100.00 | 100.00 | 100.00 |
| 10 | 100.00 | 100.00 | 100.00 | — | — | — | — | — | — |

**Fig 2.**
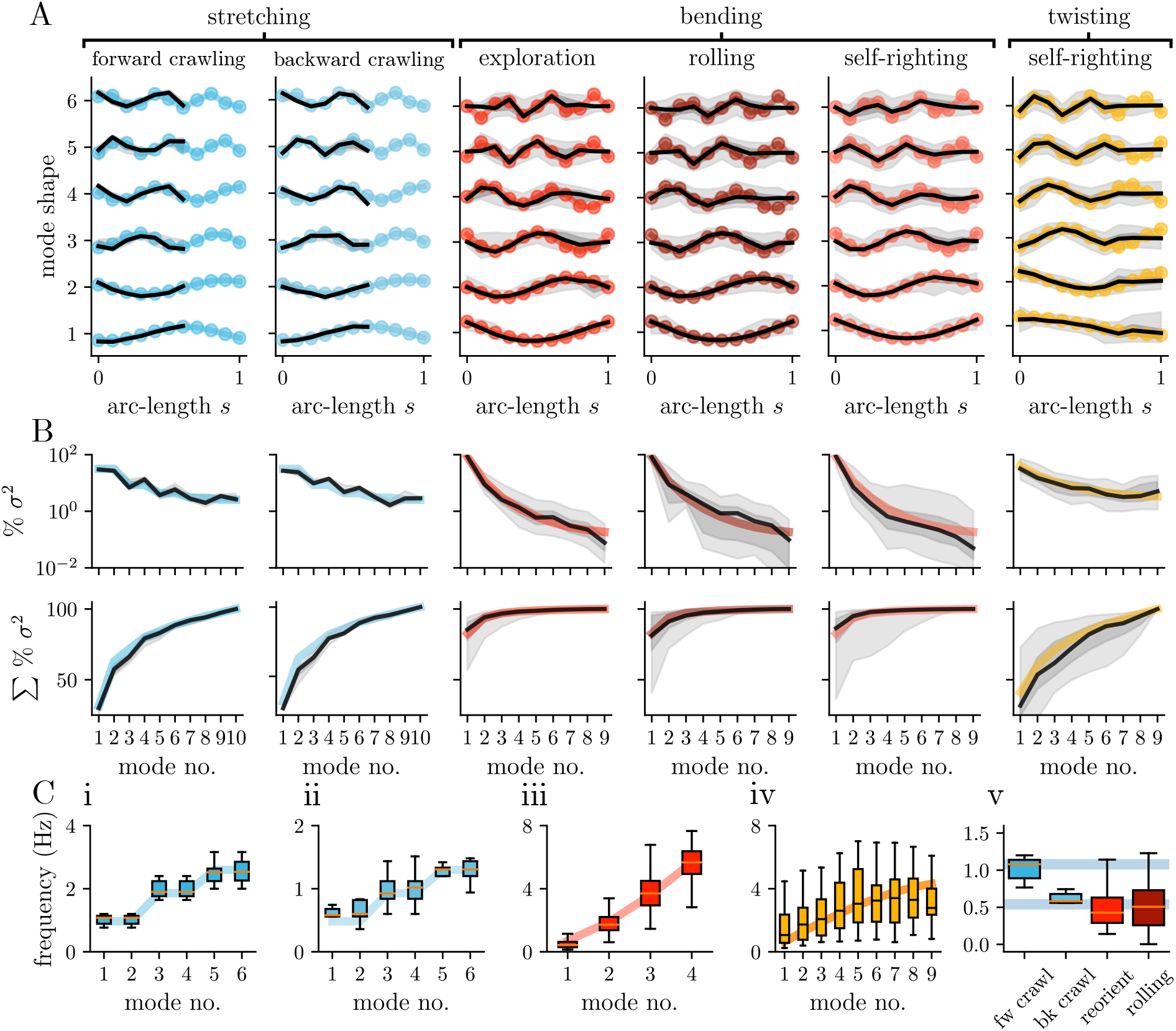
Modal analysis of the effective theory predicts the principal components and dimensionality of larval behaviour. A) First six mode shapes predicted by our effective theory (coloured lines) compared to principal components of stretching (extracted from recordings of forward and backward peristaltic crawling behaviour), bending (extracted from recordings of unbiased substrate exploration, rolling, and self-righting behaviour) and twisting (extracted from recordings of self-righting behaviour) deformations. PCA was performed on individual larvae. Median principal component shown in black, interquartile range in dark grey, and 2nd–98th percentile range in light grey. Downward deflection corresponds to segmental compression (stretching modes), right-handed bending (bending modes) or clockwise rotation (twisting modes). Mode shape predictions are parameter-free. B) Predicted proportion of variance explained by each theoretical mode shape (coloured lines) compared to observed proportion of variance along each theoretical mode during real behaviour (median in black, interquartle range in dark grey, 2nd–98th percentile range in light grey). Data are shown for individual modes (top) and cumulatively (bottom). Larval behaviour is low-dimensional in that a large proportion of variance can be explained by the first few longest wavelength modes in each behaviour, as predicted by our theory. C) The frequency relationships observed between stretching modes during forward (i) and backward (ii) crawling, bending modes during unbiased exploration (iii), and twisting modes during self-righting (iii) compared to the fit by our model (boxplots show real data; coloured lines show model fit; fit obtained by tuning the three free parameters *ω*_*s*_, *ω*_*b*_, *ω*_*t*_ using nonlinear least squares). (v) Frequencies of the first stretching mode during forward/backward crawling (fw/bk crawl) and the first “C-bending” mode during unbiased exploration (reorient) compared to the angular velocity of rolling. Our theory correctly predicts that the angular velocity of rolling should match the frequency of C-bending during exploration (*p* = 0.23, Mann-Whitney U-test) but incorrectly predicts that the frequency of forward and backward crawling should match (*p* = 0.009).

### 1.4 Principal components analysis

We compare the mode shapes predicted by our effective theory to those obtained via PCA of real larval behaviour in Figure 2. There is a strikingly good agreement between theory and experiment across all of the deformation modes and larval behaviours we studied: axial compression/expansion observed during forward and backward peristaltic crawling, mediolateral transverse bending during unbiased substrate exploration behaviour, mediolateral–dorsoventral transverse bending observed during self-righting and rolling behaviours, and twisting observed during self-righting (Figure 2A).

Furthermore, the proportion of variance explained by each of our predicted modes is surprisingly well explained by equipartition of energy, with substantial variance explained by the first few long-wavelength modes (Figure 2B and Table 1). Indeed, our equilibrium model predicts that 80.10% of variance in stretching should be explained by the first four axial modes; on average, these modes explained 79.34% and 78.70 of axial variance during forward and backward crawling, respectively. Similarly, we predict that 82.67% of variance in transverse bending should be explained by the first transverse mode alone; on average, this “C-bending” mode explained 85.72%, 82.01%, and 90.22% of transverse variance during unbiased exploration, rolling, and self-righting behaviours, respectively. Meanwhile, we predict that 71.23% of variance in twisting should be explained by the first three torsional modes; on average, these modes explained 70.19% of torsional variance during self-righting. Since our model is able to predict the proportion of variance along each mode, it is also able to directly predict the “PCA dimension” of larval behaviours, i.e. the number of modes required to explain some threshold proportion of variance (usually set to between 70–90% of variance) [49]. We list our predicted dimensions along with those computed from real data in Table 2, where we have used linear interpolation to predict the (fractional) dimension required to explain 70% or 90% of variance, providing lower and upper bounds on the PCA dimension. Again, there is good agreement between theory and experiment, with a maximum error between our predicted lower and upper dimensions and the data-estimated dimensions of 0.65. Thus, we believe our model provides a plausible explanation for the observed low-dimensionality of larval behaviour: shorter wavelength modes suffer from Boltzmann suppression. This means that all modes have the same *energy* on average, but this amount of energy produces much larger *amplitude* excursions in the long-wavelength modes than in their short-wavelength counterparts. As a result, observations of larval behaviour are dominated by the long-wavelength modes.

**Table 2.**
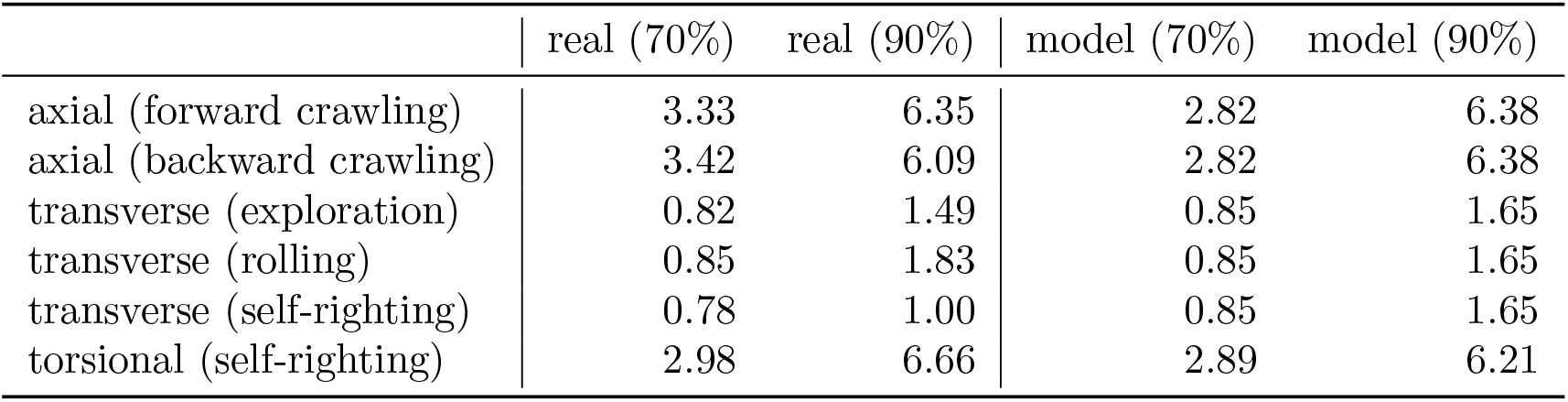
Number of modes required to explain 70–90% of variance during different larval behaviours, compared to model prediction. These percentages represent the range of commonly used thresholds when estimating the dimensionality of a system via PCA [49]. Reported values were computed by linear interpolation of data in Table 1.

|  | real (70%) | real (90%) | model (70%) | model (90%) |
| --- | --- | --- | --- | --- |
| axial (forward crawling) | 3.33 | 6.35 | 2.82 | 6.38 |
| axial (backward crawling) | 3.42 | 6.09 | 2.82 | 6.38 |
| transverse (exploration) | 0.82 | 1.49 | 0.85 | 1.65 |
| transverse (rolling) | 0.85 | 1.83 | 0.85 | 1.65 |
| transverse (self-righting) | 0.78 | 1.00 | 0.85 | 1.65 |
| torsional (self-righting) | 2.98 | 6.66 | 2.89 | 6.21 |

One of the striking results of our analysis is that the eigenmaggot shapes are largely conserved between unbiased behaviour and rolling behaviour – our theory also gives a strong explanation of why this must be the case: these shapes are encoded in the effective physics of the midline. This, would otherwise be, highly non-intuitive. For instance, it may be surprising that similar proportions of transverse variance are explained by C-bending during unbiased behaviour (85.72%) and rolling (82.01%), despite this type of deformation being more visually obvious during rolling. However, our equilibrium effective theory tells us that this should be true simply because equipartition of energy amongst the modes should generically lead to a large amount of variance being explained by C-bending.

The predictions so far have been independent of the particular values of our models’ parameters. We can now fix some of these parameters before moving forward, by considering the timescales of axial, transverse, and torsional deformations during larval behaviour. In particular, we have tuned the three free natural frequency parameters of our theory (which determine the relative timescales of the axial, transverse, and torsional deformations, as well as the overall absolute timescale of our model) to fit the observed frequencies for each mode. Indeed, we were able to find qualitatively good fits to these spectra (Figure 2C), although we were required to *separately* fit the spectrum of forward and backward peristaltic crawling as the overall timescales of these behaviours differ from one another [21,50]. Explaining this discrepancy will require future modification of our effective theory.

We note that we have determined the average frequency ratio between the first mediolateral bending mode during unbiased behaviour and the first axial mode during forward crawling to be ≈ 0.4 (reported value is the ratio of median frequencies for the two behaviours). Our low-energy effective theory is able to fit, but not explain, this frequency ratio due to the lack of energetic interactions between axial and transverse modes. However, in Appendix D we extend our theory to a higher energy regime by including a prototypical nonlinear axial-transverse interaction (based on the geometric/kinematic nonlinearity in [19]); this interaction may lead to transfer of energy between the axial and transverse fields via a 2: 1 parametric resonance, thus predicting an axial-transverse frequency ratio of ≈ 0.5, close to our measured value.

At this point, we are left with the temperature *β* as the only free parameter of the theory; this is joined by an additional parameter in the following section.

### 1.5 Non-Gaussian Statistics and Degeneracy

Although the Gaussian statistics of our theory successfully predict mode shapes and the proportion of variance along each modal coordinate during real larval behaviour, they will in general fail to adequately describe the probability distributions of these coordinates, which can be strongly non-Gaussian (Figure 3; Anderson-Darling normality test, *p* < 0.01 for all distributions of modal coordinates).

**Fig 3.**
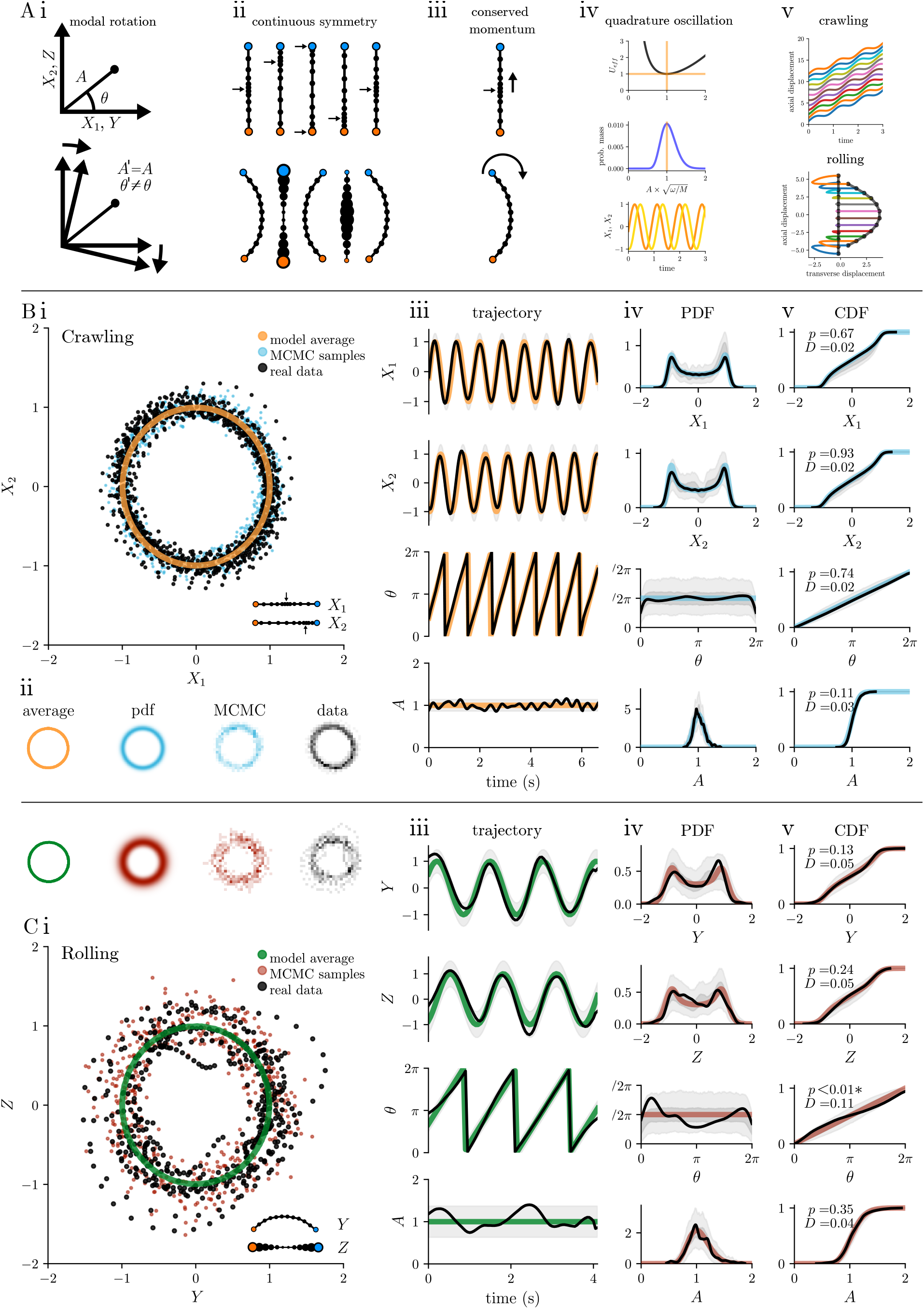
Similar physics underlies crawling and rolling behaviours. Ai) (top) A pair of modes with equal (degenerate) frequency *ω* can be described in terms of their respective modal coordinates (e.g. we use *X*_1_, *X*_2_ for first pair of axial modes, and *Y, Z* for first mediolateral and first dorsoventral bending mode) or in terms of their amplitude *A* and phase *θ*. (bottom) Since our effective energy depends only on *A* and not *θ*, our model is symmetric under continuous rotations of the degenerate modes. Aii) This rotation corresponds to moving the peak of axial compression along the body, considering the head and tail as rigidly linked via the visceral piston [21] (rotation of *X*_1_, *X*_2_ plane; top) or to rotating the direction of a C-bend around the body axis (rotation of *Y, Z* plane, bottom) Aiii) By Noether’s theorem [47] the rotational invariance is linked to a conserved mechanical invariant (generalised momentum *M*) corresponding to the momentum of a compression wave propagating along the body (top) or a C-bend rotating around the body axis (bottom). Aiv) Incorporating the conserved momentum into the theory as a constraint gives rise to an effective potential energy (*U*_*eff*_, top) for the amplitude *A* [52]. The associated Boltzmann statistics for *A* can then be computed (middle); the most probable (average) trajectory minimizes the effective potential (orange lines) and has angular momentum *M*, corresponding to oscillation of the degenerate modes in quadrature (bottom) with amplitude 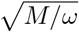 and frequency *ω*. Av) The quadrature modal oscillations correspond to peristaltic compression waves (top; oscillation in *X*_1_, *X*_2_) or rolling (bottom; oscillation in *Y, Z*). B, C) Fit of the model average trajectory and Boltzmann statistics to real data for forward crawling (B) and rolling (C); amplitudes were normalised so that 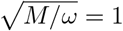. i) Average trajectory (unit circle), real data (black), and a matched number of Markov chain Monte Carlo (MCMC) samples from the Boltzmann distribution (coloured circles) in the degenerate mode plane. ii) separate plots of the data in (i), including the analytical Boltzmann probability density function (pdf) iii) Representative trajectories of modal coordinates, amplitude, and phase over time (black) compared to average trajectory (coloured line) iv) Kernel density estimate (KDE, black) of the pdf of each quantity in (iii), computed over all trials and compared to predicted pdf (coloured line), with two predicted error bounds corresponding to either 2nd–98th percentile for MCMC samples drawn from the predicted pdf (light grey; see Methods) or interquartile range from the same MCMC samples (dark grey). v) cumulative distribution function (cdf) for each quantity in (iv) (same colorscheme;), with results of Kolmogorov-Smirnov comparisons between the predicted cdf and the corresponding empirical distribution functions.

This discrepancy might be corrected by perturbatively modifying the Hamiltonian of our theory in order to bring the predicted statistics closer to those we observe. Indeed, such perturbative modifications to a theory are a cornerstone of the effective theory approach [35, 36]. However, in our case we can make some progress simply by looking closer at the symmetries of our Hamiltonian, without modification. In particular, we note that our choice of periodic boundary conditions for the axial field, representing the visceral piston [21] – which mechanically couples the head and tail extremities – introduces an effective rotational symmetry into our theory. We similarly introduced a rotational symmetry though our choice to set the mediolateral and dorsoventral frequency parameters to be equal, representing the approximately circular cross-section of the animal. Both symmetries manifest as degeneracies in the eigenvalue spectrum of our Hamiltonian – in other words, the axial stretching/compression modes appear in pairs with identical frequency, and similarly for every mediolateral bending mode there is a corresponding dorsoventral mode with the same frequency.

In our theory, due to Noether’s theorem [51], the symmetries mentioned above are associated with mechanical invariants: quantities which, like energy, do not change during the evolution of a closed system. The derivation is presented below.

We start from the Hamiltonian for a pair of modes with degenerate frequency

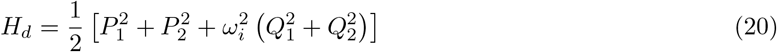

here, *ω*_*i*_ is the common frequency of the two modes with coordinates *Q*_1_, *Q*_2_ and conjugate momenta *P*_1_, *P*_2_. To make the rotational symmetry of this Hamiltonian manifest, we take a canonical transformation to polar coordinates in the *Q*_1_ – *Q*_2_ plane, by setting *Q*_1_ = *A* cos *ϕ, Q*_2_ = *A* sin *ϕ*, where *A* is the “amplitude”, or distance from the origin, in the plane, and *ϕ* is the “phase”. In terms of these new coordinates we obtain the Hamiltonian

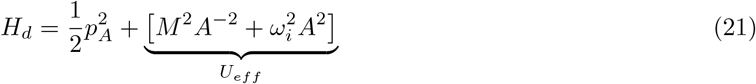

where *p*_*A*_ is the canonical momentum associated with the amplitude coordinate *A*, and *M* is the canonical momentum associated with the phase coordinate *ϕ*, corresponding to the angular momentum in the *Q*_1_ – *Q*_2_ plane. Note that *ϕ* does not appear in the Hamiltonian. This signifies that there is no energetic “cost” associated with rotations in the *Q*_1_ – *Q*_2_ plane – this is our internal rotational symmetry. As a consequence, the angular momentum *M* is a mechanical invariant. This is a straightforward consequence of Hamilton’s equation for the dynamics of *M*,

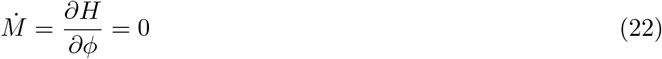

Since the time derivative of *M* is zero, *M* itself must be conserved – it stays constant over time. This canonical momentum can therefore be treated as a parameter in the Hamiltonian, defining an effective potential energy for the amplitude coordinate (we have labelled this *U*_*eff*_ in Equation 21). We formally derive the maximum entropy distribution subject to a fixed value of *M* and a fixed ensemble average of the Hamiltonian *H*_*d*_ in Appendix I, finding

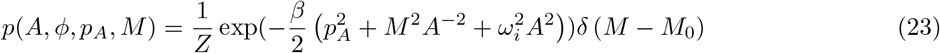

This can be interpreted as the usual Boltzmann statistics for *H*_*d*_ multiplied by an additional Dirac delta function which constrains *M* to take a fixed value *M*_0_. We calculate the partition function to be

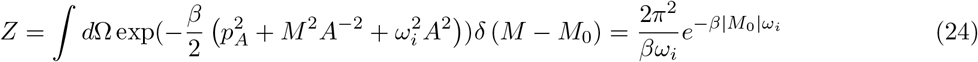

where the measure *d*∂ represents integration over phase space. Note that the most probable behaviour of the system can be obtained from a saddle point approximation to the partition function integral, corresponding to minimisation of the Hamiltonian *H*_*d*_ subject to *M* = *M*_0_. This minimisation is straightforward, and tells us that the amplitude takes a fixed value 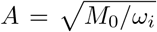, which is nonzero for all *M*_0_ = 0, and with zero associated momentum *p*_*A*_ = 0. Thus, the most probable behaviour consists of a circular trajectory in the *Q*_1_ – *Q*_2_ plane, in which our original modal coordinates execute sinusoidal oscillations in quadrature (i.e. with a 90° relative phase shift).

### 1.6 Analyses of larval peristalsis and rolling behaviours

We now turn our attention to the peristaltic crawling and rolling behaviours of the larva. Both behaviours involve a continuous, ongoing, seemingly out-of-equilibrium, motion of the body. In peristalsis, a wave of segmental compression and expansion passes along the body. In forward crawling, this wave travels from the tail of the animal, along its anteroposterior axis, until it reaches the head. At this point the head, tail, and internal viscera move in synchrony [21], and a new wave is initiated at the tail. In backward crawling the behaviour is similar, except the wave travels from head to tail and propagates at around half the speed of its forward–propagating counterparts (Figure 2). In both cases, the wave has a single “peak” of maximal compression, but is spread across several segments, so that it appears largely as a broad sinusoid of compression and expansion (recall that the first two axial modes alone explain around 55% of variance during these behaviours, Table 1). In rolling, the larva enters a C-bend configuration (corresponding to the first bending mode of our model – recall that this mode explains around 80% of variance during the behaviour, Table 1). The C-bend stays in the plane of the substrate while the body rotates, so that the C-bend propagates around the body axis in a clockwise or counterclockwise direction. For clockwise rolling, the C-bend might first be directed to the right of the body, then ventrally, then to the left, then dorsally, then back to the right; for counterclockwise rolling the sequence is reversed.

To the eye, rolling and peristalsis behaviours could scarcely look more different. However, owing to the similar underlying rotational symmetry of the axial and transverse fields in our model, and the resulting degeneracy of the axial and transverse spectra, both behaviours should share be described in mode space by the degenerate statistics that we derived in the previous section, differing only in the values of the invariant *M*_0_ and the inverse temperature *β*.

In Figure 3 we plot experimental estimates for the first pair of degenerate modes during peristalsis (first two axial compression modes, labelled *X*_1_, *X*_2_, Figure 3B) and during rolling (first mediolateral and first dorsoventral bending mode, labelled *Y, Z*, Figure 3C), respectively, alongside a fit of our model to this data. To perform our fit, we first eliminated *M*_0_ by normalising the modes by the location of the peak of the amplitude distribution (this peak lies at 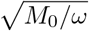 in our model; Since the free parameter *ω* was fixed in the previous section, normalisation is equivalent to fixing *M*_0_). We then tuned *β* to fit the normalised amplitude distribution using nonlinear least squares. We also plot our estimate of the (average) behavioural trajectory both in the normalised mode space, where it corresponds to the unit circle, and as time series for the individual modes, their amplitude *A*, and their phase *ϕ*. For both behaviours, the fit by our model is qualitatively very good. Indeed, we find only one statistically significant difference between the predicted and observed kinematic distributions, for the phase *ϕ* during rolling (KS test, *p* < 0.01, Figure 3Cv; all other comparisons have *p* ≥ 0.11). This distribution is predicted to be uniform, but shows a peak near 0° (which corresponds to pure mediolateral C-bending), possibly reflecting the transitions to/from rolling at the initial and final portions of our recordings.

We also note that our estimate of the average behavioural trajectory provides a novel experimental test of our theory of rolling. In particular, our estimated trajectory tells us that the average angular velocity with which the C-bend rotates about the body axis should simply be given by the frequency of the first transverse mode *ω*_*b*,1_. Indeed, this parameter should set the dominant timescale of the first transverse mode across all behaviours. Thus we predict that the average frequency of C-bending during unbiased behaviour should match the average angular velocity of rolling. Indeed, we find no statistically significant difference between our estimates of these two quantities (Figure 2; two-sided Mann-Whitney U-test, *p* = 0.4).

### 1.7 Study of bending deformations during larval substrate exploration

Next we consider the statistics of bending deformations during substrate exploration. In this case, we can expect to obtain a correct description when *M*_0_ = 0 for the transverse modes, since this means that bending deformations will remain plane [41, 42] rather than rotationally propagating around the midline as in rolling behaviour.

Setting *M*_0_ = 0 and assuming all bending occurs in the mediolateral plane, we can write the distribution for the *i*’th mediolateral modal coordinate *Y*_*i*_ as

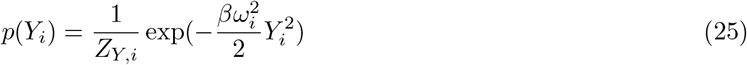

with the configurational partition function

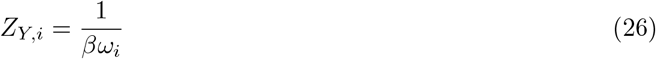

Thus in the case *M*_0_ = 0 the predicted modal distributions are again zero-centred Gaussians with variances 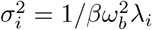, in line with our earlier results in which the momenta *M* were neglected.

We fit the free parameter *β* to fit the variances 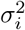 of the all the observed modal distributions during unbiased behaviour. In Figure 4 we plot the first five modal distributions on axes normalised by our predicted *σ*_*i*_; with this normalisation, our model predicts that all of the modal distributions should correspond to the standardised Gaussian distribution with zero mean and unit variance. There is again a qualitatively good agreement between theory and experiment. The observed distributions are approximately zero-centred (absolute mean |*m*| ≤ 0.028 for all distributions), have variances which are well fit by our model (see Figure 2B), and have low skew (absolute skew |*S*| ≤ 0.52 for all distributions), but have heavier tails than our predicted Gaussian distributions (Fisher’s kurtosis *K* ≥ 3.9 for all distributions, compared to *K* = 0 for normal distribution), leading to statistically significant differences for all comparisons between our model’s predicted bending mode distributions and the observed distributions, although the Kolmogorov-Smirnov statistical distance between predicted and observed distributions remains small (KS test distance *D* ≤ 0.09, *p* < 0.01 for all comparisons). The inability of our effective theory to explain the tails of the transverse bending distributions during unbiased behaviour is unsurprising given our restriction to the relevant low-energy physics, since large amplitude bends should be associated with higher elastic energies. Given the qualitatively good fit between our predicted Gaussian statistics and the observed bending mode distributions, we believe our effective theory should still be seen as a useful starting point for the development of higher energy theories, and can hopefully still provide a useful explanatory model.

**Fig 4.**
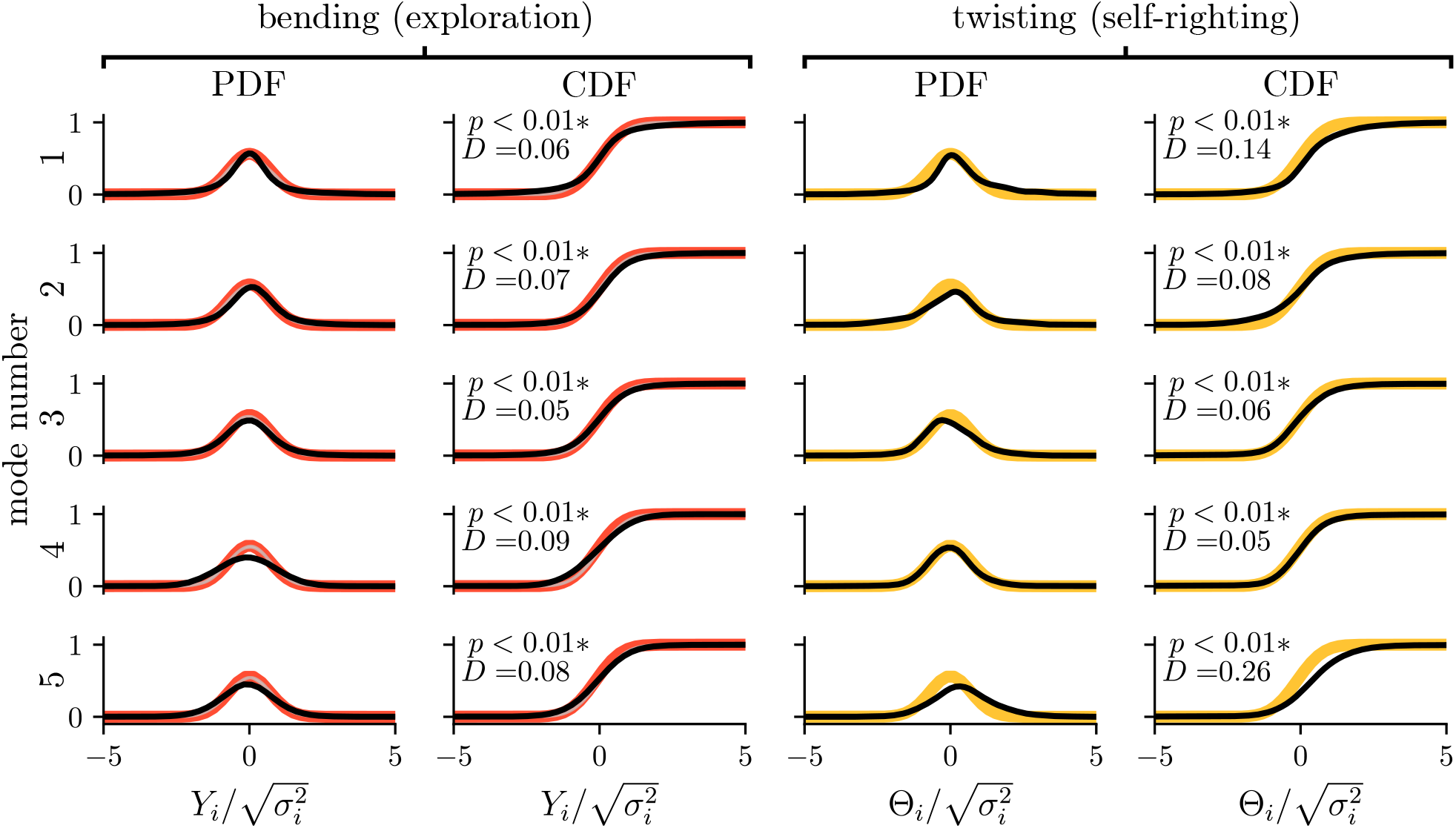
Approximately Gaussian statistics of bending and twisting deformations. Kernel density estimate (black) of the probability density function (pdf) and cumulative density function (cdf) for the first five bending modes *Y*_*i*_ (during unbiased exploration, left) and the first five twisting modes Θ_*i*_ (during a mixture of unbiased exploration and self-righting, right) with Kolmogorov-Smirnov comparisons to the distributions predicted by our model. Each mode was normalised by the mode-specific standard deviation 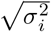, which is fixed by our model up to an overall scaling by *β*. We determined this parameter by fitting the variance of the modes using nonlinear least squares. After normalisation, our model predicts each modal coordinate to have a zero-centred Gaussian distribution with unit variance (red and yellow lines). Although our model closely predicts the observed distributions (KS distance *D* ≤ 0.09 for all bending distributions and *D* ≤ 0.08 for 3 of 5 twisting distributions), all comparisons reached statistical significance (*p* < 0.01; see text for discussion).

### 1.8 Statistics of twisting

The degenerate frequencies of axial and transverse modes resulted from underlying continuous symmetries – periodic boundary conditions for the axial modes, and radial symmetry about the midline for the transverse modes. The torsional modes possess no such underlying symmetry, and so are non-degenerate. In this case, our model predicts that the torsional modes should have Gaussian statistics, with the relative variances of the torsional modes fixed by the eigenvalues of the second difference matrix with free boundary conditions, and the overall variance determined by the inverse temperature *β* of the Boltzmann distribution (Section 1.3).

As for the bending deformations during unbiased exploration (see previous section), our Gaussian model provides a qualitatively good approximation to the distributions of the torsional mode amplitudes (which we were able to observe during a mixture of both self-righting and unbiased exploration behaviour; Figure 4). However, all of the measured distributions show statistically significant differences when formally compared to the Gaussian model predictions (KS test *D* = 0.16 ± 0.10, *p* < 0.01 for all comparisons). This is unsurprising, given that we expect large torsional deformations to lie outside the domain of validity of our low-energy theory. We believe our *qualitative* Gaussian approximation should still be seen as a useful starting point for the development of higher energy theories, and should still provide a useful explanatory model.

### 1.9 Model-driven experimental characterisation of larval self-righting behaviour

Finally, we turn our attention to the self-righting (SR) behaviour of the larva. This behaviour can be observed by experimentally preparing larvae in an “upside-down” configuration with their dorsal surface in contact with the substrate, after which the larva is able to turn itself so that its ventral surface is in contact with the substrate, and the animals’ normal spontaneous behaviour resumes [16]. At present, there is no widely accepted theoretical, mechanistic explanation for how the larva accomplishes self-righting. An obvious explanation would be that the larva simply uses the same mechanism that it uses to roll (as described previously in this paper), which we will call the rolling-based SR model. However, an alternative explanation has been posited [16, 25, 26], which suggests that during self-righting, and unlike during rolling behaviour, the larva twists its body so as to attach its mouth hooks to the substrate, before rotating the rest of its segments to bring the body fully into alignment with the substrate (Figure 5A). We will call this alternative explanation the twist-based SR model. Armed with our theory of 3-dimensional larval movement, we are well placed to formalise these explanations and assess exactly how self-righting is accomplished. Since we have already explored rolling behaviour earlier in this paper, we will now focus on the twist-based SR model.

**Fig 5.**
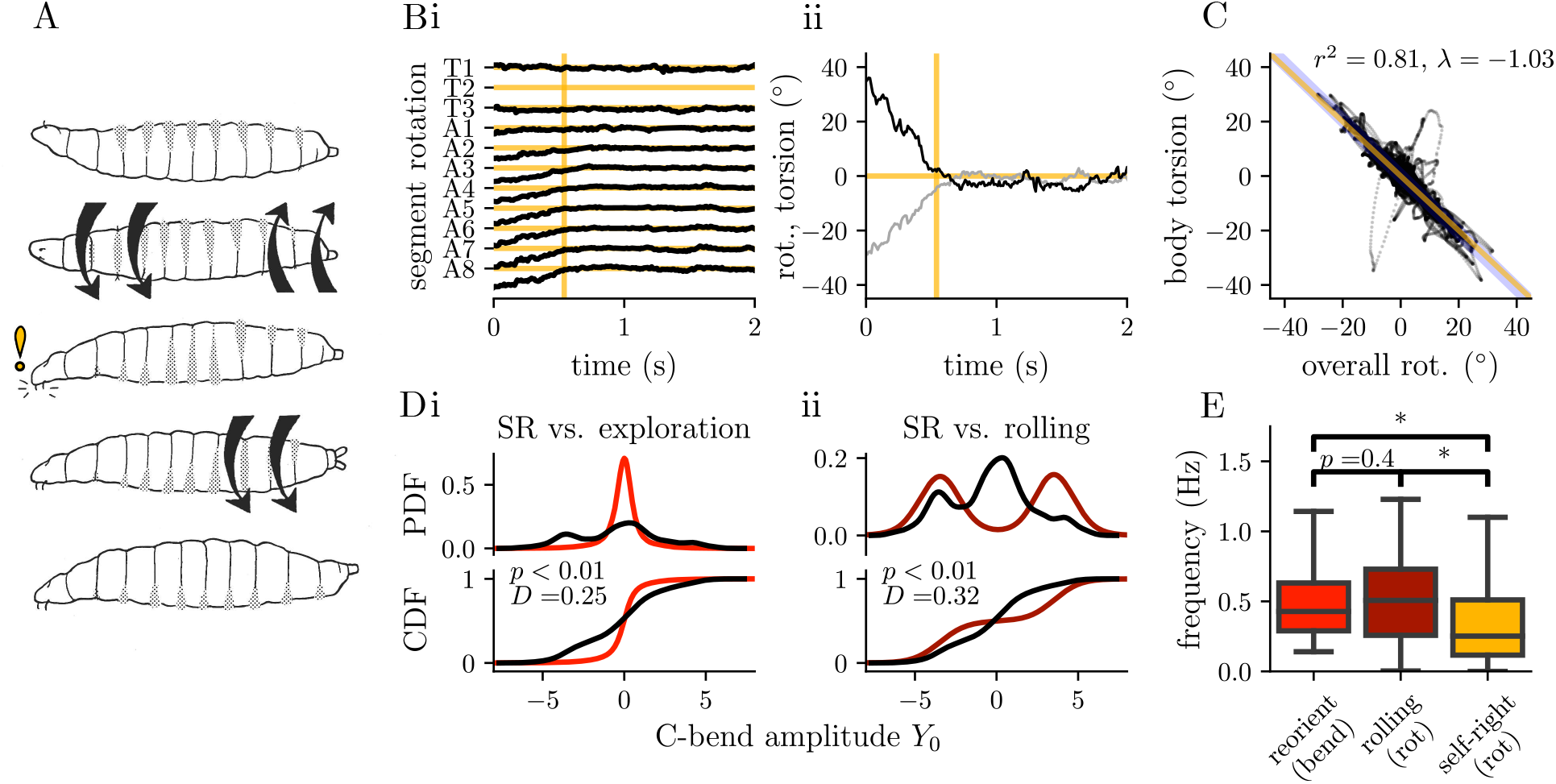
Model-driven investigation of self-righting behaviour suggests the larva rights itself using twisting deformations. A) In the twist-based model of self-righting (SR), the larva begins in an “upside-down” configuration (top), twists along its length to attach its mouth hooks to the substrate (middle) and then un-twists to align its body with the substrate (bottom). Alternatively, the larva may self-right via rolling (roll-based SR, see text). Bi) Representative trajectory showing the rotation angle of all body segments (apart from T2) about the body axis during the final portion of SR behaviour; note that the thoracic segments (T1, T3) remain in fixed alignment with the substrate throughout, while abdominal segments (A1–A8) rotate to align with them, until eventually the body is completely aligned with the substrate (yellow lines). Bii) overall rotation of the body (black) and estimated torsion along the body (grey, torsion estimated using first two torsional modes) for the trajectory shown in (Bi). C) overall rotation and torsion along the body are strongly negatively correlated (Pearson’s *r*^2^ = 0.81), with a slope near that predicted by the twist-based SR model (measured *λ* = −1.03, blue, vs predicted *λ* = −1, yellow), strongly suggesting Di, Dii) Kernel density estimates of the pdf and cdf for the first bending mode, measured in the plane of the substrate, during SR (black), substrate exploration (bright red), and rolling (dark red). The distribution during SR is closer to that for exploration (Kolmogorov-Smirnov distance *D* = 0.25) than that for rolling (Kolmogorov-Smirnov distance *D* = 0.32) and all distributions differ significantly (Kolmogorov-Smirnov test *p* < 0.01 for all comparisons) E) Furthermore, the angular velocity during rolling (dark red) matches the frequency of C-bending during unbiased exploration (bright red; *p* = 0.23, Mann-Whitney U test) while the angular velocity during self-righting differs from both (yellow, *p* < 0.01, Mann-Whitney U test), suggesting that self-righting is not driven by bending as in the roll-based SR model.

We first observe that the passive mechanical equilibrium of our effective theory, i.e. the minimum energy configuration, has all segments aligned with one another, since in this configuration there is no energy stored in the twisting or bending of body segments relative to one another. Thus, after attachment of the mouth hooks to the substrate, the minimum energy configuration (which corresponds to the most probable configuration due to the negative exponential weighting of energy in the Boltzmann distribution) will consist of *all* segments being aligned with the substrate. In other words, attachment of the mouth hooks to the substrate should be sufficient to produce full self-righting, via subsequent “passive” relaxation towards equilibrium.

We can formulate several experimental tests of this twist-based SR model. First, this model suggests that the head and mouth parts of the larva should not rotate relative to the substrate during SR, while the other parts of the body should rotate into alignment with the head and mouth parts. Indeed, this is exactly what we observe (Figure 5B), in contradiction to the rolling-based SR model, which would require the head and mouth parts to rotate along with the rest of the body during SR.

Second, the overall (average) rotation of the larva relative to the substrate, captured by the zero-frequency torsional mode, should be strongly correlated with the higher torsional modes. This follows because the attachment of the mouth hooks to the substrate introduces a constraint on the torsional modes, which is exactly what allows overall rotation of the larva to be driven by torsional deformations during self-righting. In particular, the following identities must hold

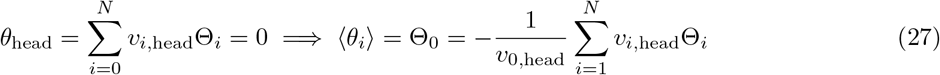

where *θ*_head_ is the rotation of the head, *v*_*i*,head_ is the element of the *i*’th torsional mode shape at the head, and Θ_*i*_ is the *i*’th torsional modal coordinate (with *i* = 0 corresponding to the zero-frequency overall rotation mode). The overall rotation of the body at any instant is given by the average over segments ⟨*θ*_*i*_⟩, which corresponds exactly with the zero-frequency mode Θ_0_. This relationship can be interpreted as a simple restatement of the fact that the overall rotation is proportional to the torsional deformation in the twist-based SR model. As our model explains, *>* 55% of the variance in torsional deformations is accounted for by the first and second torsional modes alone. We should therefore have the approximation

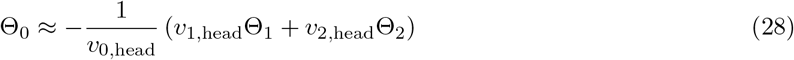

Experimentally, we have indeed observed a very strong correlation between the overall rotation and the torsion accounted for by the first two torsional modes (representative trajectory Figure 5Bii, correlation plot Figure 5C; Pearson’s *r*^2^ = 0.81) with a slope parameter very close to the predicted relationship (observed *λ* = *−*1.03 vs. ideal *λ* = *−*1). Although there is a statistically significant difference between the observed correlation slope and that predicted (*p* < 0.01 two-sided Wald test under null hypothesis that *λ* = *−*1), this is to be expected given the approximate nature of Equation 28. Note again that this observation is in contradiction to the rolling-based SR model, which would require there to be no such constrained relationships between the torsional modes; if there was such a constraint, the larva would produce a very large, clearly observable torsion during rolling as Θ_0_ increases approximately linearly during rolling, sometimes for several complete rotations (Figure 3). Unexpectedly, the correlation between overall rotation and torsion remains present even for very small overall rotations, i.e. rotation and torsion are correlated even when the larva is righted. This suggests that twist-based SR may be an ongoing process that occurs *during* other behaviours.

Thirdly, the rolling-based SR model suggests that self-righting should be driven by similar bending deformations as those seen during rolling behaviour, rather than the twisting deformations predicted by the twist-based SR model. Therefore, the rolling-based SR model predicts that the angular velocity associated with overall rotation during self-righting, i.e. 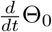, should be equal to the average frequency of C-bending during unbiased substrate exploration behaviour. This relationship holds during rolling, with the angular velocity of rolling closely matching the average frequency of C-bending during exploration (median = 0.46 Hz for rolling; median = 0.43 Hz for exploration; Mann-Whitney U-test *p* = 0.23; Figure 5E, see Section 1.6). However, during self-righting behaviour, the rate of overall rotation is lower than both the C-bending frequency during exploration, and the rate of overall rotation during rolling (median = 0.25 Hz for self-righting; Mann-Whitney U-test *p* < 0.01 for both comparisons). This result is again in favour of the twist-based rather than rolling-based model of self-righting.

Finally, we directly compare the distribution of C-bend amplitudes during self-righting, rolling, and unbiased substrate exploration. We find that the C-bend amplitude distribution during self-righting is closer to that observed during unbiased exploration than that observed during rolling (KS distance, *D* = 0.25 SR vs unbiased exploration, *D* = 0.32 SR vs rolling; Figure 5D). However, all three distributions differ from one another (KS test *p* < 0.01 for all comparisons): the C-bend distribution is unimodal and approximately zero-centred Gaussian, the rolling distribution is bimodal (with peaks corresponding to large-amplitude left-handed or right-handed C-bending) and zero-centred, and the self-righting distribution is bimodal and not zero-centred. Qualitatively, one of the peaks in the SR distribution appears to coincide with the peak of the C-bend distribution for unbiased behaviour, and the other appears to coincide with one of the peaks of the rolling distribution. Thus, although C-bending does not appear to play the same *role* during SR as it does during rolling (since the first three experimental tests point to SR being driven by twisting rather than bending), large-amplitude C-bending does appear to be a clear feature of SR. It is unclear why this should be the case. One possibility is that C-bending during SR is a result of twist-bend coupling, a phenomenon that occurs due to the geometrically nonlinear rod kinematics that become apparent at large deformations (higher energy scales) [53], and which is not accounted for in our (linear) low-energy theory. Alternatively, this observation may be due simply to our inability to measure segmental rotations greater than ∼ 90° due to occlusion of the anatomical landmarks used during tracking (see Methods), so that we are only able to measure the final stages of SR; perhaps the initial stages of SR consist of rotation driven by C-bending, as in rolling behaviour, and the larva only attaches its mouth hooks once it is close to being righted. This would explain why we observe a large-amplitude peak in the C-bend distribution despite the fact that overall rotation appears to be driven by twisting rather than bending. The resolution of this issue is unfortunately beyond the scope of this paper; further progress must wait until either a higher energy theory is developed or experimental limits are extended.

## 2 Discussion

In this paper we develop an effective theory for the study of movement in small animals. In particular, our work presents a model of the low-energy physics of the Drosophila larval AP axis (midline), and experimental demonstration of this model’s ability to predict real behavioural data. Our model is the first to describe the fully 3-dimensional motions of the larval midline, including stretching along, bending of, and twisting about the body axis. We believe it sets the stage for development of similar theories for the behaviour of a large class of small animals with “slender” or “rod-like” body morphologies, including the invertebrate nematode worm *C. elegans* and the vertebrate zebrafish *D. rerio*. Our theory may also be extended to the behaviour of larger animals such as snakes.

### The relation of theory to experimental observations

Our effective theory makes several predictions, many of which we have directly tested. For instance, our theory makes parameter-free predictions of the principal components of stretching, bending, and twisting, as well as the proportion of variance explained by each component. These predictions closely match the results of PCA applied to real larval deformations during a range of behaviours including forward/backward peristaltic crawling, unbiased substrate exploration, rolling, and self-righting (Figure 2). Since our effective theory correctly predicts the proportion of variance associated with each mode, it may provide a simple explanation for the low-dimensionality of larval behaviour, via Boltzmann suppression of short wavelength modes. In this view, all of the deformation modes have the same average energy (due to equipartition), but because the longer wavelength modes are “softer” than the “stiff” short wavelength modes, this energy causes larger fluctuations in the longer wavelength modes. Thus, observations of behaviour are dominated by the first few long wavelength modes.

Our theory is also able to predict low-dimensional average trajectories for the larva’s rhythmic rolling and peristalsis behaviours (Figure 3). It is also able to very closely predict the stretching and bending mode distributions during these behaviours, and provides a good first approximation to the bending and twisting mode distributions observed during substrate exploration and self-righting behaviours (Figure 3 and 4). As a test of our model’s utility, we were also able to use it to investigate the mechanism underlying self-righting behaviour; our analysis in terms of our model’s modes supported a view of self-righting being driven by twisting deformations, rather than the bending deformations that drive rolling behaviour.

Given our model’s explicit focus on the low-energy physics of the larva, we expect it to break down at the higher energy scales governing short-wavelength and large amplitude deformations of the body. Indeed, although our model provides a qualitatively very good fit to the mode shapes extracted from recordings of real animal behaviour, the fit is certainly better for the dominant long-wavelength modes than for their short-wavelength counterparts (Figure 2). Furthermore, although we successfully predict the *variance* structure of bending and twisting deformations, and our predicted Gaussian statistics for the corresponding distributions in mode space are therefore qualitatively very good, our theory does not account for higher-order statistics. We are thus unable to account for the heavy tails of these distributions, which represent relatively high-energy, large-amplitude bending and twisting deformations. Given our theory’s success at low energies, however, we are hopeful that it may provide a useful starting point for a perturbative approach to understanding the higher energy regime [35].

The model we propose raises several interesting conceptual questions – in particular, it remains unclear, first, why our assumption of statistical equilibrium provides such a good description of larval behaviour; and, secondly, why our theory works despite containing no detailed description of the larva’s neuromuscular system, environment, or detailed features of the (presumably nonlinear) biomechanics. We consider these two questions in detail below.

### Statistical equilibrium and its power to explain larval behaviour

We constructed our effective theory of the larval midline using an approach borrowed from statistical field theory [36], in which emphasis is placed on the symmetry and stability requirements that strongly constrain the theoretical description of a physical system at low energies. The theory that we obtained by applying this method essentially predicts that the larval midline should be governed by the statistical mechanics of linear elasticity [43], similar to the theoretical description of polymer mechanics via wormlike chain models [45]. This similarity follows simply from the observation that the larval midline and long polymers possess similar symmetries; both systems can be efficiently described by “infinitely thin” curves in space, with invariance under overall translations and rotations as well as reflection invariance. These symmetries lead the low-energy descriptions of these disparate systems to coincide. However, whereas the statistical equilibrium of polymer models usually has a relatively straightforward explanation in terms of essentially “random” collisions of the polymer with surrounding solvent molecules [45], here the success of the statistical equilibrium hypothesis in predicting features of larval behaviour is highly surprising and requires further consideration.

Crucially, our use of equilibrium statistical mechanics necessarily presupposes some mechanism that is not contained directly in our effective theory. This should be clear since the effective theory is described by a quadratic Hamiltonian, which should give rise to linear deterministic dynamics [51], and so our theory contains no mechanism by which trajectories may “wander” or “mix” in the phase space [54]; in other words, the deterministic dynamics of our theory cannot explain its predicted statistics without modification. This is actually a fundamental issue at the foundations of equilibrium statistical mechanics, and not specific to our theory alone, since for very many systems to which equilibrium statistics are applied it is either not possible to prove that the underlying dynamics of the system are capable of driving the system to equilibrium, or it is possible to show that the dynamics are outright incapable of this feat (as in the case of our effective theory) [40, 54].

In some cases, effective equilibrium statistics may arise even though the underlying system is known to be out of equilibrium [55,56]; presumably this must occur in the case of the larva, whose body mechanics experience a strong flow of energy from the musculature. Indeed, it is interesting to consider how equilibrium statistics may arise in larval behaviour. One argument is that small nonlinear modifications to our effective theory should generically lead to energetic coupling between modes; it is quite possible that such coupling may lead to chaotic dynamics, providing a mechanism by which the modes may thermalise. Our previous work on the planar mechanics of unbiased exploration [19] suggests that beyond the low energy limit we have considered here, the mechanics of bending and stretching of the midline should indeed become energetically coupled, leading to chaotic motion. However, it is currently unclear whether this planar model is sufficiently ergodic to explain the appearance of equilibrium statistics. Recent work in the nematode *C elegans* has experimentally demonstrated the presence of a symmetric Lyapunov spectrum for the animal’s dynamics, strongly suggestive of deterministic chaos arising due to (damped, driven) Hamiltonian mechanics [13, 57]; if such a spectrum is present in larval behaviour, it may go some way towards explaining the animals’ behavioural statistics. Alternatively, the presence of noisy forcing may also be a factor in explaining thermalisation (we comment on this below).

### Accurate prediction of larval behaviour without a detailed description of the neuromuscular system

We note that our theory is able to explain features of larval behaviour without including a detailed description of the animals’ nervous system. We argue that it is able to do so by capturing the essential combined *effects* of the body physics and the musculature. That is, the free parameters in our theory capture the role of both body mechanics and neuromuscular forcing. The first three of our parameters control the natural frequency scales for stretching, bending, and torsional deformations. It is clear that the timescale of peristaltic crawling can be experimentally influenced by perturbing either the central nervous system [58–61] or feedback from peripheral sensory neurons [62,63], suggesting that the effect of both the nervous system and body mechanics should be captured by our phenomenological axial natural frequency parameter.

The temperature parameters in our theory govern the average energy of, and the strength of fluctuations in larval deformations. We have commented above on how this parameter may represent the effects of unmodelled dynamics in the theory, including the effect of *mechanical* coupling between stretching and bending deformations, which may lead to deterministic chaotic dynamics [19]. However, it is also clear that this parameter must include the role of the neuromuscular system and the environment. Indeed, maintaining an average energy in the larva must require a constant input of energy from the musculature to balance frictional losses to the environment. Furthermore, this parameter may reflect the role of *neuromuscular noise*. Indeed, inclusion of noisy forcing into the linear deterministic dynamics of our current effective theory could give rise to a Langevin dynamics capable of explaining the appearance of equilibrium statistics [40, 64].

The final parameters in our theory, the momenta *M*, which we associate with the deformations during crawling and rolling, govern the (average) amplitude of axial compressions during peristaltic crawling and transverse C-bending during rolling. Again, although we derived these momenta as the mechanical invariants related to our theory’s underlying rotational symmetries via Noether’s theorem, the maintenance of these momenta in the presence of environmental perturbations and noise is likely to be a crucial effect of the neuromuscular system.

Although our effective theory does not directly contain a description of the larval nervous system, this does not mean that it has nothing to say about the role of the neuromuscular system. Indeed, by providing a description in which the phenomenological parameters support multiple mechanistic interpretations (see above), our theory is able to contribute to advancing our fundamental understanding of movement control, by focusing effort on teasing apart the contributions due to intrinsic properties of body mechanics, from those produced via neural actions and sensory processes. For example, the outcome of recent genetic screens aimed at mapping genes (e.g. microRNAs) affecting larval movement [25, 26, 65] can now be pursued considering effects of gene modulation and control on body mechanics, as well as, on neural development, function and control. In addition, we can also envisage projections of our work into neural aspects of robotic design and development: a theoretical understanding of what body mechanics can achieve *per se*, can help define the minimal requirements needed to effectively command robotic movement.

More generally, our theory suggests that considerations of the effective physics of the body can go a long way to explaining complex and diverse behaviours. In this context, the role of the nervous system may be seen as being more modulatory in nature – shifting the global phenomenological parameters of the system – rather than representing a precise micro-management system acting to control the exact trajectories of the animal body during behaviour.

Given the simplicity of our starting assumptions we believe our effective theory may be readily applicable to studying the behaviour of other animals with similar morphologies to the Drosophila larva. This includes several other important “model species” such as the nematode worm *C. elegans*, and the zebrafish *D. rerio*, thus opening the way to the investigation of both, invertebrate and vertebrate systems.

## 3 Methods

### 3.1 Self-righting behaviour assays

Self-righting (SR) assays were conducted as done previously [16] on 1st instar wild-type larvae (w1118), separately to our experiments on unbiased behaviour and rolling. Briefly, parental lines were raised at 25°*C* in collection cages bearing apple juice-based medium agar plates, supplemented with yeast paste. From these plates, stage 17 embryos [66] were collected and transferred to a fresh plate. Next, freshly-hatched first instar larvae were transferred again on 0.9% agarose plates. After one minute of acclimatisation, we conducted self-righting assays by gently rolling over the larvae with a small brush pen to an upside-down position. Wild type larvae normally take approximately 6–7 seconds to rectify their position (self-right). SR sequences were recorded at 10 fps with a Basler ace acA800-510*µ*m (CMOS) monochrome USB 3.0 Camera mounted on the Leica MZ75 stereomicroscope.

### 3.2 Peristalsis assay

The data we used to quantify peristalsis behaviour was previously collected and published in [50]. Videos of forward and backward crawling were collected at 30 fps. The lengths of abdominal segments were measured at the left and right sides of the body; we used the mean of left and right measurements to estimate the midline segment length prior to further analysis (see below).

### 3.3 Rolling and unbiased exploration behaviour assays

#### 3.3.1 Fly strains

For behavioural assay of unbiased behaviour and rolling, we used R69F06-GAL4 and a control line with no GAL4 expression, w;;attp2, from the Rubin GAL4/LexA collection [67, 68]. These were crossed to UAS-CsChrimson::mVenus [69] for optogenetic activation of rolling behaviour. For mechanical nociceptive stimulation behavior experiment, CantonS were used. All flies are raised at 25°C for 4 days – 3rd instar larvae before wandering stage animals are used for behaviour experiment.

#### 3.3.2 Behavioural apparatus

The apparatus comprises a video camera (FLIR GS3-U3-51S5M-C camera, 2048 × 2048), for monitoring larvae, a light illuminator (LED 624 nm) for optogenetic activation, and a computer, similar to [23]. Recordings were captured at 30 fps and were controlled through the Multi-Worm Tracker (MWT) software (http://sourceforge.net/projects/mwt) [70], whilst control of the hardware module was controlled through the Stimulus Control Module (SCM) software. For mechanical stimulation, video was captured 30 fps using FLIR software.

#### 3.3.3 Behavioural experiments

Embryos were collected for 24 hours at 25°C. Foraging third instar larvae were used for all experiments. Larvae were raised in the dark at 25°C for 4 days on fly food containing trans-retinal (Sigma, R2500), at a concentration of 200 *µ*M. Before an experiment, larvae were separated from food by suspension in 15% sucrose and then washed with water. Larvae were dried, then transferred to the centre of a 25 × 25 cm transparent plastic, square plate covered in a layer of 2% agar gel. Between 20–60 larvae were transferred to the plate for any given recording. Optogenetic stimulation was delivered in two 15 second bouts at an irradiance of 600 *µ*W/cm2, with a 30 second interval between bouts. Experiments were run under infrared light to avoid tonic activation of UAS-CsChrimson::mVenus. For mechanical nociceptive stimulation, single larvae were placed on 2% agar gel. After larvae started crawling, they were pinched by forceps (Dumont, N5) to evoke rolling behavior.

#### 3.3.4 Behaviour quantification

For calculation of larval midlines during exploration and rolling, behavioural recordings were captured with the Multi-worm Tracker (MWT) software http://sourceforge.net/projects/mwt [70]. MWT returns a 2D contour (outline) for each larva tracked, at 30fps. From these contours, we computed the larval midline using the choreography package (bundled with MWT) [70]. The estimated midline is given by a set of equally spaced points, allowing characterisation of transverse bending but not axial compression/expansion.

Larval midlines and contours were used to score bouts of rolling behaviour, as described elsewhere [71]. Larvae that were tracked for fewer than 5 seconds, or travelled less than one body length in distance, were rejected. To determine angular velocity of rolling behaviour, videos were manually scored using the Fiji software (NIH) [72]. First, we scored the angle of larval rotation to the nearest 60 degree interval (0, 60, 120, 180, 240, 300), where 0 degrees represented dorsal-side up, and 180 degrees represented dorsal-side down. The trachea and posterior spiracles were used as landmarks to determine the degree of rotation around the anteroposterior axis. The duration of a roll was demarcated by the first and last frame in which the larva presented movement perpendicular to it’s body axis. The mean angular velocity of rolling was calculated as the total angle rotated divided by the duration of roll.

For calculation of larval deformations during self-righting, DeepLabCut (version 2.2b5) was used. Using a standard DeepLabCut pipeline [73], at least 20 distinct frames were extracted per studied video using DLC’s K-means clustering. Up to 44 features were manually labelled per frame according to their visibility. These included points consisting of a Contour (Thoracic Segments [T1-3 left and right], Abdominal Segments [A1-8 left and right]), Mouth Hooks and a Tracheal System (Abdominal Segments [A1-8 Trachea left and right] and Posterior Spiracles). These manually annotated frames served as a training dataset fed into the ResNet-50 provided by DLC. The ResNet-50 was trained for 1,000,000 iterations via batch processing across NVIDIA V100 GPUs. All video frames were then analysed using this trained network to produce estimated poses. We then estimated the midline using the mean of the left and right contour markers, and estimated a tracheal centroid by taking the mean of the left and right trachea markers.

We estimated the segmental rotation angles *θ*_*i*_ between the substrate surface normal and the local sagittal plane as *θ*_*i*_ = tan^−1^(*d*_*i*_*/r*_*i*_), where *d*_*i*_ is the measured distance between the midline and the tracheal centroid at the *i*^t^*th* segment, and *r*_*i*_ is the radius of the body at that segment. We estimated the segmental diameter (and hence radius) as being equal to the measured width of the larva in our camera projection, at each video frame.

### 3.4 Comparison of experiment to theory

For unbiased exploration, rolling, and self-righting behaviour, postural principal components (“eigenmaggots”) were extracted from the midline data using the singular value decomposition (SVD)-based principal components analysis (PCA) provided by the `scikit-learn` python machine learning module [74]. Prior to PCA, we removed the overall translational and rotational degrees of freedom of the animal by computing the angles between segments [75]. For self-righting behaviour, we additionally performed PCA on the estimated abdominal rotation angles.

For peristalsis behaviour, principal components were extracted from measurements of abdominal segment lengths during forward and backward crawling (data previously published in [50]) using the same SVD-based PCA decomposition as for unbiased behaviour and rolling behaviour. Overall translation was not measured in this dataset so did not need to be removed prior to PCA.

For comparison to our theory, the experimental data was also projected onto the modal basis of our effective theory. We projected the abdominal segment length data onto the eigenvectors **V**_2,*c*_ of the circulant second difference matrix **D**_2,*c*_, which is identical to a spatial discrete Fourier transform basis (Appendix E), to obtain estimates of the axial modal coordinates *X*_*i*_. We also normalised to correct for the truncation of the axial modal basis (since only abdominal segments were measured). In particular, we modelled the axial displacements as arising from a measurement process

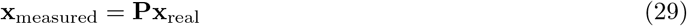

where **P** is the matrix which projects the “full” vector of axial displacements **x**_real_ onto the subspace of abdominal displacements (i.e. **P** models our lack of thoracic data). We wish to estimate the amplitudes **X** of the axial modes. These mode amplitudes are defined via **x** = **V**_2,*c*_*X*. Thus, we have

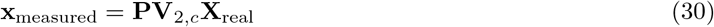

Ideally, we would like to invert this expression in order to find **X**_real_ given **x**_measured_. We can use the orthonormality of the eigenbasis to write 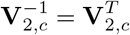 (orthonormality follows from the fact that the circulant second difference matrix is real and symmetric). However, the projection matrix is singular (as are all projection matrices). We therefore use the Moore-Penrose pseudoinverse projection **P**^T^ = **P**^+^ to write an initial estimate of the mode amplitudes using the truncated basis vectors 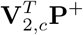,

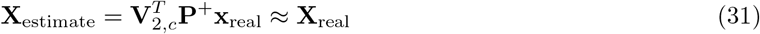

Using our expression for **x**_real_ in terms of **X**_real_ lets us write

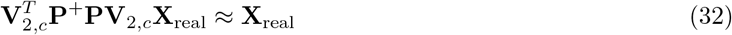

Which tells us our approximation will be improved by bringing the transformation on the left closer to the identity transformation. We cannot rotate our basis vectors to orthogonalise them (since our purpose is to estimate the deflections along the *particular* basis vectors in **V**_2,*c*_). However, we can at least bring our transformation closer to an identity by normalising our new basis. To see this, imagine we have a pure deflection *X*_*i*_ along a single mode vector *v*_*i*_. Then our transformation would give the approximation

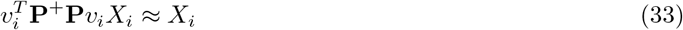

which can clearly be made exact by normalising by the scalar 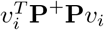. Writing all such normalising factors in the matrix 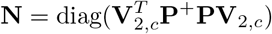 gives the improved approximation

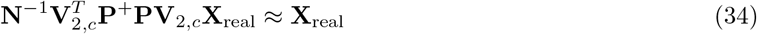

Finally, we again use our expression for **X**_real_ in terms of **x**_measured_ (along with the weak-inverse property of the pseudoinverse **P**^+^**PP**^+^ = **P**^+^) to write our estimate of the mode amplitudes as

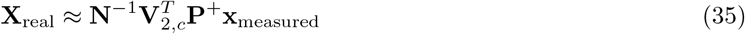

In addition to applying this transformation, for figure 3 we further applied a brickwall (0.5–1.5Hz) bandpass filter to the first pair of axial modes to remove noise and artifacts associated with the truncation.

The estimated abdominal and thoracic rotation angles ***θ*** were projected onto the eigenvectors of the free-boundary second difference matrix *D*_2,*f*_. Since we were unable to measure the rotation of the second thoracic segment T2, we estimated the torsional mode amplitudes using a similar method as for the axial modes, applying the transformation

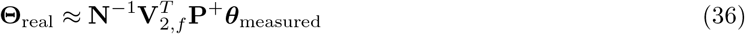

where now 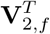is the eigenbasis of the free-boundary second difference matrix and **P** is the projectionmatrix modelling our lack of measurements for T2.

The midline data from unbiased exploration, rolling, and self-righting behaviour experiments was projected onto the eigenvectors of the free-boundary fourth difference matrix *D*_4,*f*_. We estimated the eigenvectors numerically using the numpy python numerics module [76] as the resulting eigenvectors are more accurate than our analytical asymptotic approximations. To remove the overall translational and rotational degrees of freedom we performed this projection using the angles between midline segments, as for PCA (above), by similarly transforming the eigenvectors into the angle space. Unbiased behaviour occurs mainly in the plane of the substrate with little rolling, so we use the projection to estimate the mediolateral transverse modal coordinates *Y*_*i*_ during this behaviour. By contrast, the larva continuously rotates during rolling so that our top–down view allows us to effectively estimate the modal amplitude in the mediolateral–dorsoventral plane, 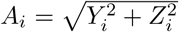 rather than the individual mediolateral (*Y*_*i*_) or dorsoventral (*Z*_*i*_) modal coordinates.

To estimate the mean modal frequencies during behaviour we first estimated the power spectral density (PSD) of each modal coordinate, for each individual larva tested, using the Welch spectrogram averaging method, implemented in the scipy python scientific computing module [77]. Since peristalsis is a highly rhythmic behaviour, the estimated PSDs are tightly peaked [50]. We therefore estimated the mean modal frequencies during this behaviour using the strongest peak in the PSD. Unbiased behaviour is aperiodic, displaying a “spread” in the PSD estimates rather than cleanly localised peaks [34]. We therefore estimated the mean modal frequencies during this behaviour using the spectral centroid, i.e. the weighted average of the PSD. The higher modes are associated with a very low variance, and for the fifth transverse mode and above the spectrum drops towards the uniform PSD expected of white noise. In this case the spectral centroid stays close to the mid-point of the spectrum; for our sampling rate of 30 Hz, the Nyquist frequency is 15 Hz and the spectral centroid of white noise is 7.5 Hz. We therefore attempted to fit only the frequencies of the first four transverse modes.

Unless otherwise noted above all statistical analyses and tests were performed using the scipy statistics submodule [77].

## Author roles and acknowledgements

Jane Loveless developed the theoretical aspects of this work, conducted data analysis for comparisons between experiment and theory, and wrote the bulk of the manuscript. Alastair Garner conducted experiments on rolling and unbiased behaviour. Abdul Raouf Issa conducted experiments on self-righting, hand-labelled the resulting data for input to DeepLabCut, and aided in experimental design for our self-righting assays. Ruairí J. V. Roberts hand-labelled self-righting data, performed DeepLabCut-based analysis of self-righting data, and aided in experimental design for our self-righting assays. Barbara Webb provided guidance on the development of the theory and aided with editing the manuscript. Lucia L. Prieto-Godino supervised the analysis of self-righting data via DeepLabCut and aided in experimental design for our self-righting assays. Tomoko Ohyama supervised collection of experimental data on rolling and unbiased behaviour, conducted experiments on rolling and unbiased behaviour, and aided with editing the manuscript. Claudio R. Alonso promoted the project, supervised the collection of experimental data (self-righting), provided guidance, and contributed to the writing and editing of the manuscript.

Data for forward/backward peristalsis was provided by Stefan Pulver. This data was previously published in [50].

Illustration of self-righting behaviour in Figure 5 was produced by Eleanor Savage.

We thank Greg Stephens, Tosif Ahamed, Antonio Carlos Costa, Matthieu Louis, Kostantinos Lagogiannis, and James Fraser for helpful discussions during the development of this project.

This research was supported by: a Wellcome Trust Investigator Award (CRA) (Ref: 098410/Z/12/Z), a Medical Research Council Project Grant (CRA) (Ref: MR/S011609/1), a Natural Sciences and Engineering Research Council of Canada Discovery Grant (TO) (Ref:04781), a Canadian Institutes of Health Research Project grant (TO) (Ref: 153030), a Fonds de Recherche Nature et Technologies Quebec New Investigator Grant (TO) (Ref: 255237), a Canada Foundation for Innovation John R. Evans Leaders fund (TO) (Ref: 36533), a Boehringer Ingelheim Fonds PhD Fellowship (RJVR), a Francis Crick Institute Grant (LLPG) (Ref: FC001594), and a European Research Council ERC StG “EvoNeuroCircuit” (LLPG) (Ref: 802531).

## A Detailed construction of the effective theory

In the main text, we summarised the construction of an effective Hamiltonian *H* for the axial, transverse, and torsional deformation fields *x, y, z, θ* of the larval midline. In this section we work through this construction in greater detail. Rather than starting directly from the Hamiltonian we will instead start from it’s Legendre transform, the Lagrangian

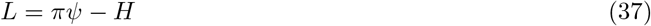

where *ψ* = [*x, y, z, t*]^*T*^ is the vector of deformation fields and *π* is the conjugate canonical momentum vector. Assuming locality, we can write the Lagrangian as an integral *L* = ∫*dsℒ* of a Lagrangian density. This Lagrangian density is the Legendre transform of the Hamiltonian density in the main text, and it can be written in terms of the fields and their derivatives at a point in space and time (*s, t*)as

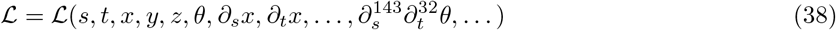

As is common in field theory, we will occasionally refer to this Lagrangian density simply as the Lagrangian, despite the fact that this term technically refers to the integral of the density. Given a Lagrangian, the conservative dynamics of the deformation fields can be obtained from the Euler-Lagrange field equations

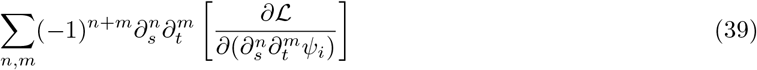

with *ψ*_*i*_ *∈ ψ*. Assuming the Lagrangian is analytic, we can perform a Taylor expansion and write

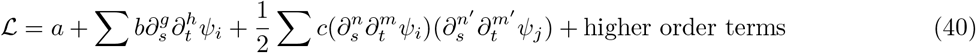

Here we have adopted the convention that the zero’th order derivative is the identity, i.e. *∂* ^0^*ψ*_*i*_ = *ψ*_*i*_ and summations are over all non-negative integer-order derivatives of the fields; indices on the constants *b, c* have been dropped (there is a unique constant for each summand).

We can discard the constant *a* since it will not appear in the Euler-Lagrange equations and therefore does not effect the physics. Similarly, any linear terms (contained in the first summation) with *g >* 0 or *h >* 0 will not contribute to the Euler-Lagrange equations and can also be discarded, leaving only linear terms of the form *b*_*i*_*ψ*_*i*_. Furthermore, only quadratic terms where the sum *n* + *m* + *n*′ + *m*′ is even will contribute to the Euler-Lagrange equations (as can be checked by the interested reader).

We now apply our first symmetry requirement: that the Lagrangian be invariant under a reflection of each of the fields *ψ*_*i*_ → *− ψ*_*i*_ individually. This removes the remaining linear terms, and constrains *ψ*_*i*_ = *ψ*_*j*_ in the quadratic terms, i.e. to quadratic order the deformation fields are now completely decoupled. This lets us write the Lagrangian as a sum of field-specific quadratic Lagrangians plus an interaction part,

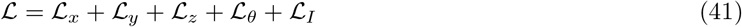

where each of the first four *ℒ*_*ϕ*_ contains only the quadratic terms for the field *ϕ ∈ {x, y, z, θ}*, i.e.

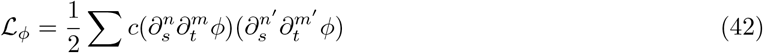

while *ℒ*_*I*_ contains all higher order terms and interactions. We will now focus our attention on the quadratic Lagrangians, while continuing to use *ϕ* as a generic field symbol to denote any of the specific fields *x, y, z, θ* in our theory. Applying our second symmetry requirement: invariance under overall translations of the fields *ϕ* → *ϕ* + Δ*ϕ*. This eliminates the zero’th order derivative term where *n* = *m* = *n*′ = *m*′ = 0, corresponding to *ϕ*^2^, and we can also safely eliminate any other term containing a zero’th order derivative (since all such terms are completely equivalent to terms without a zero’th order derivative on going to the Euler-Lagrange equation). In other words, the only remaining terms must be of a minimum order with *n* + *m ≥* 1 and *n*′ + *m*′ *≥* 1.

Turning our attention to the Lagrangians for the transverse *y* and *z* fields, we can now apply our third symmetry requirement: invariance under overall rotations of the fields. For an infinitesimal rotation through angle *δ* about the center of the midline (*s* = 1*/*2), this takes the form *ϕ* → *ϕ*+(*s−*1*/*2)*δ*. This condition rules out the term *c*(*∂*_*s*_*ϕ*)^2^, and thus all remaining terms must be of minimum order 2*m* + *n ≥* 2 and 2*m*′ + *n*′ *≥* 2 in *ℒ*_*y*_ and *ℒ*_*z*_.

Now we focus our attention on the low-energy (long-wavelength, low-frequency, low-amplitude) physics of the midline, by using a form of dimensional analysis inspired by the renormalisation group [36, 40]. We first note that by focusing on low energies we are interested primarily in small deformations of the larva, characterised by small amplitudes of the fields and their derivatives, and we can therefore discard the interaction Lagrangian *ℒ* _*I*_ entirely (technically some terms in the interaction could be required to counteract instabilities of the fields [36], but we will soon see that we are lucky in this respect).

We proceed by introducing a rescaling of space *s* = *σs*′ and time *t* = *τt*′ into our theory, along with a compensatory rescaling of the field amplitude *ϕ* = *εϕ*′. For *τ >* 1, *σ >* 1 we are essentially “zooming out” and viewing the larva on longer length and time scales. As we do so, the constants in our Lagrangians change, since on rescaling we have

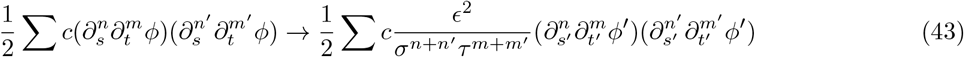

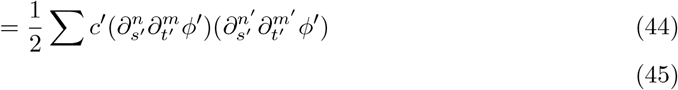

with the ratio of the constants before and after the rescaling transformation given by

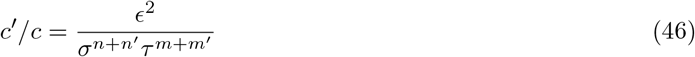

We can use our amplitude rescaling parameter *ε* to “fix” a certain set of constants within the Lagrangian, allowing us to observe how the other constants grow or shrink relative to the fixed set. Since we have discarded the interaction Lagrangian, we can focus on each of our field-specific quadratic Lagrangians as though each governed a completely separate subsystem. We start with the Lagrangians for *x* and *θ*, for which we determined the minimum order terms to be given by *n*+*m* = 1 and *n*′ +*m*′ = 1 on the basis of translation and reflection invariance. There are three unique terms satisfying these conditions, namely the terms containing (*∂*_*s*_*ϕ*)^2^, (*∂*_*t*_*ϕ*)^2^ and (*∂*_*s*_*ϕ*)(*∂*_*t*_*ϕ*). Fixing the coefficient of the first term gives *c*′*/c* = 1 = *ε*^2^*/σ*^2^ = *⇒ ε* = *σ*, while fixing the coefficient of the second term gives *c*′*/c* = 1 = *ε*^2^ = *τ* ^2^ = *⇒ ε* = *τ* = *σ*. Substituting these into the coefficient of the third term gives *c*′*/c* = *ε*^2^*/στ* = 1 so that this term neither grows nor shrinks relative to the first two. What about the higher order terms? We can substitute *ε* = *τ* = *σ* into the growth equation (Equation 46) to find the relationship

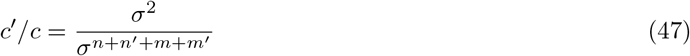

This tells us that in *ℒ*_*x*_ and *ℒ*_*θ*_, all terms satisfying 2 < *m* + *m*′ + *n* + *n*′, i.e. all terms above the lowest order, will shrink on rescaling. We can follow the same argument for the *y* and *z* fields. In this case we determined the minimum order terms to be given by the condition 2 = 2*m* + *n* and 2 = 2*m*′ + *n*′. There are just two unique terms satisfying these conditions, namely the terms containing 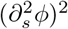 and (*∂*_*t*_*ϕ*)^2^. Fixing these terms during rescaling gives *ϵ* = *τ* = *σ*^2^. The higher order terms in *ℒ*_*y*_ and *ℒ*_*z*_ then change on rescaling as

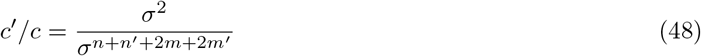

which tells us that all terms satisfying 2 < *n* + *n*′ + 2*m* + 2*m*′ will shrink on rescaling; again, this corresponds to all terms containing derivatives higher than the minimal order.

On repeated rescalings, most of the constants in our theory flow towards zero exponentially fast, and only the minimal order derivatives maintain non-zero coefficients. At the fixed point, we are therefore left with the Lagrangians

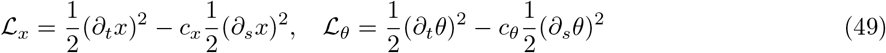

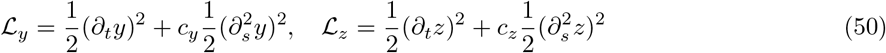

where we have removed the coefficient of the first term in each Lagrangian through an appropriate redefinition of the fields, leaving 4 free parameters. The first pair of Lagrangians, governing the axial deformations *x* and the torsional deformations *θ*, are recognisable as giving linear wave mechanics. Meanwhile, the second pair, governing transverse bending deformations *y, z*, are recognisable as giving Euler-Bernoulli beam mechanics. Thus, our derivation tells us that the low-energy physics of the larva should be dominated by the classical linear elasticity theory of deformable rods [52]. We note that our theory is thus strongly reminiscent of the phenomenological elastic models of that have been used with great success in understanding mechanical properties of DNA and other polymers [40, 44–46], with the key difference being the inclusion of kinetic energy, owing to the larva’s relatively large size and our interest in the larva’s dynamics.

## B Momentum-space coarse-graining causes no flow in non-interacting theory space

In this section we provide a proof that the coarse-graining step of renormalisation causes no flow of the coupling constants within the non-interacting (Gaussian) theory space. To show this, we will start from the partition function path integral

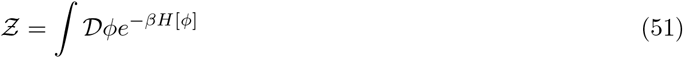

for a single scalar field *ϕ ∈ ψ* = [*x, y, z, θ*]^*T*^.

In what follows, we will perform a “momentum-space” coarse-graining. As a preliminary requirement, we must rewrite the field *ϕ* in terms of its spatial Fourier components *ϕ*_*k*_, which are labelled by their wavenumber *k* (which plays the role of momentum in quantum field theories, hence the nomenclature); intuitively, *ϕ*_*k*_ measures the magnitude of a sinusoidal excitation of the *ϕ* field with wavenumber *k*. In terms of these Fourier modes, the partition function is given by

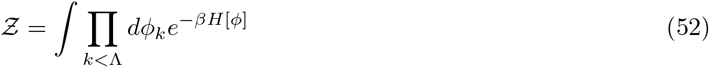

where the high wavenumber (short wavelength) cut-off Λ is imposed because we expect our long-wavelength effective theory to break down beyond some short distance scale *∼* 1*/*Λ. To enact our coarse-graining, we introduce a new cut-off Λ′ < Λ, and redefine the Fourier modes *ϕ*_*k*_, the Hamiltonian *H*, and the partition function in terms of Λ′. In particular, we split the modes *ϕ*_*k*_ into low-wavenumber and high-wavenumber modes as

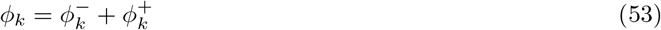

where the low-wavenumber modes are given by

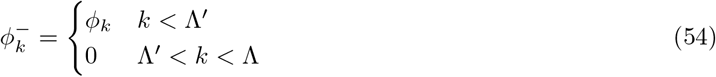

and the high-wavenumber modes are given by

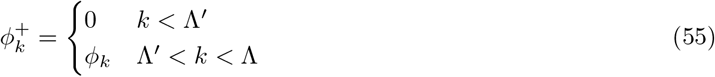

A generic Hamiltonian can then be written as

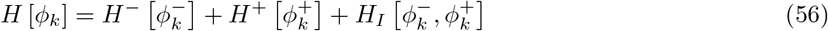

where *H*^*−*^, *H*^+^ depend only upon either the low- or high-wavenumber modes, respectively, whereas *H*_*I*_ governs interactions between the low- and high-wavenumber modes. The partition function now becomes

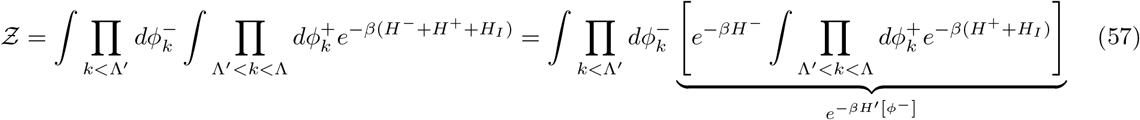

The bracketed term on the right hand side can now be used to define a new effective, coarse-grained Hamiltonian *H*′ [*ϕ*^*−*^] for the low-wavenumber modes, from which we have integrated out the dependence upon the high-wavenumber modes, as indicated by the underbrace.

In our particular case, we are interested only in the low-energy behaviour of the larval midline. We can therefore keep terms only up to quadratic order in *H* (see previous section) and neglect any higher-order contributions. In this case, the *H* must be diagonalised in Fourier space, so that *H*_*I*_ = 0 and there are no interactions between the Fourier modes. This means that the integral within the brackets above must evaluate to a constant which is independent of the low-wavenumber modes, i.e. we can simply write

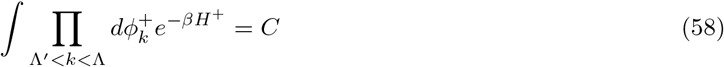

so that the coarse-grained Hamiltonian is given by the expression

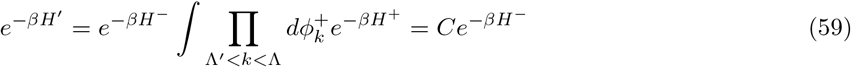

Taking the logarithm and rearranging then gives

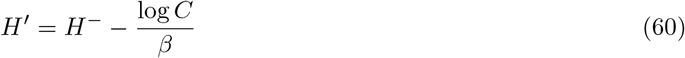

But since addition of a constant to the Hamiltonian cannot effect any physics, the new coarse-grained Hamiltonian *H*′ can be taken as being exactly equal to the original low-wavenumber Hamiltonian *H*^*−*^. Note that this means that, starting from a quadratic Hamiltonian, the coarse-graining step of renormalisation cannot cause any change in the parameters within the Hamiltonian, and all RG flow must be generated purely by the rescalings of *s, t*, and *ϕ*, all of which we accounted for in the previous section. By contrast, at higher energy scales, the presence of interactions between modes will cause *H*_*I*_ ≠ 0, so that the coarse-graining integration over the high-wavenumber modes will in general cause a change in the parameters in the low-wavenumber effective Hamiltonian.

## C Accounting for dissipative effects within Lagrangian field theory

In this section, we wish to provide an intuitive understanding for how a (fractional) quadratic Lagrangian density is able to account for the types of dissipation encountered within systems modelled by linear partial differential equations (PDEs). In particular, we will attempt to demonstrate how the general 2-dimensional scalar linear PDE of the form

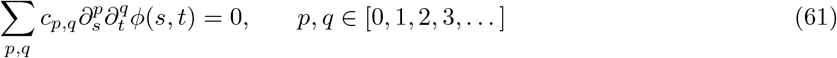

can be derived from a quadratic Lagrangian, via the Euler-Lagrange equation

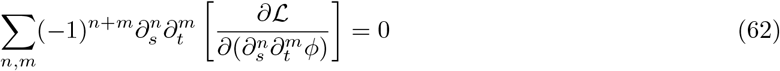

We start from the quadratic Lagrangian

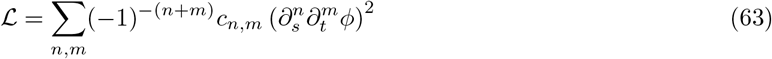

Applying the Euler-Lagrange equation then gives the PDE

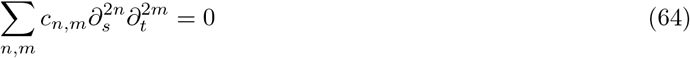

This tells us that if integer-order derivatives appear in the Lagrangian, then only even-order derivatives can appear in the Euler-Lagrange equation; this is the case usually considered, and leads to purely conservative behaviour.

However, intuitively, if we were to extend the Lagrangian to contain half-integer-order derivatives *n, m ∈* [0, 1*/*2, 1, 3*/*2, 2, 5*/*2, …], with possibly imaginary coefficients due to the factor of (*−* 1)^*−*(*n*+*m*)^, and provided the principle of least action results in a suitably generalised Euler-Lagrange equation, then the resulting dynamics should be capable of containing any integer-order derivative (i.e. derivatives of both odd and even order). In this case a simple relabelling scheme *p* = 2*n, q* = 2*m* recovers the original general 2-dimensional scalar linear PDE (Equation 61). Indeed, the Euler-Lagrange equation does generalise to this case [48].

## D Weak axial–transverse interactions and the crawling/exploration frequency ratio

We next consider the frequency relationship between the axial and transverse modes. Since the axial and transverse fields do not interact in our theory, it cannot tell us their relative frequencies. However, we can make good progress towards this aim by incorporating only a small nonlinear coupling into our theory.

Indeed, we have previously analysed the coupling between axial and mediolateral transverse degrees of freedom [19] using a simplified Hamiltonian for the axial and mediolateral transverse motions of the larva’s head

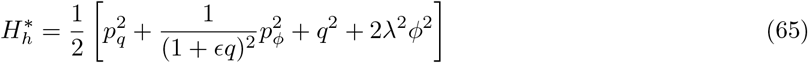

where *q* and *ϕ* are dimensionless variables characterising the axial stretch and mediolateral transverse bending angle, *p*_*q*_ and *p*_*ϕ*_ are the canonical momenta conjugate to these variables, *λ* is the axial–transverse frequency ratio, and *ε* is a global amplitude scale which multiplies the dimensionless quantities *q, ϕ* to give the real axial stretch and mediolateral bending angle of the head.

In the limit of small amplitudes the axial and transverse degrees of freedom are uncoupled, as in the effective theory we present this paper. We are interested in small coupling corrections to the uncoupled behaviour, so we expand the head Hamiltonian in a perturbation series in the amplitude parameter *ε*

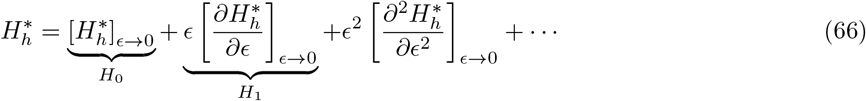

To keep our focus on the simplest model demonstrating axial-transverse coupling, we will keep only the zero-order small-amplitude term *H*_0_ and the first order correction *εH*_1_, finding

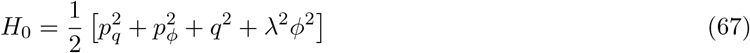

And

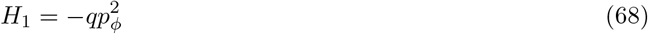

so that our intermediate amplitude Hamiltonian becomes

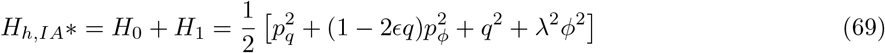

Since this coupled system is still fairly difficult to analyse, we choose to focus only on the transfer of energy from axial to transverse degrees of freedom. To do so, we will set the axial motion *q* to a prescribed function of time. We choose for this purpose *q* = cos(*ωt*), which could represent either an unperturbed axial modal vibration or a first-order approximation of the Fourier series expansion of a more complicated periodic function. Since *q* and *p*_*q*_ are now prescribed functions of time, the terms in 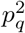 and *q*^2^ in the Hamiltonian will not effect the dynamics and can be discarded, leaving us with

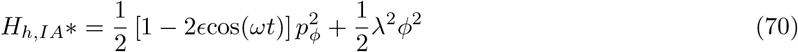

In order to simplify this Hamiltonian further, we next take a canonical transformation to new phase space coordinates given by Φ = *p*_*ϕ/λ*_, *P* = *− λϕ*, i.e. we scale and interchange the coordinates and momenta, which allows us to group all parameters in one term and interpret the axial-transverse interaction as a sinusoidal modulation of frequency

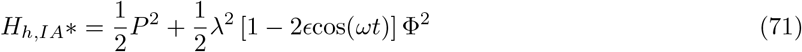

Finally we derive the intermediate amplitude transverse dynamics via the Hamilton’s equations, finding

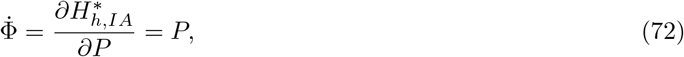

and

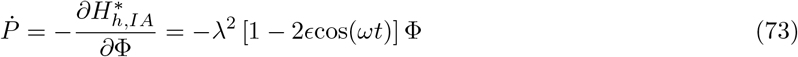

or, converting to second-order form by differentiating the first equation with respect to time and substituting into the second, we can write the dynamics in momentum space as

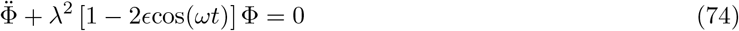

which is in the form of the Matthieu equation [78]. It is well known that this equation exhibits the phonemenon of *parametric resonance*, in which the passively stable equilibrium at Φ = 0 becomes unstable for certain values of *ε* and *ω*, giving rise to solutions which grow with time.

For infinitesimal parametric perturbations (i.e. *ε* → 0), resonance occurs when the axial forcing frequency *ω* is exactly an even multiple of the natural frequency *λ* (i.e. *ω* = 2*λ, ω* = 4*λ*, etc.). For larger perturbations (*ε >* 0), resonance occurs for a larger spread of frequencies centred on the even multiples of the natural frequency [52, 78].

In the presence of friction, larger perturbations are required to produce resonance, and the magnitude of the required perturbation grows with the forcing frequency, so that only the low-order resonances are practically accessible [52, 78]. The most readily excited resonance is therefore the 2: 1 axial-transverse resonance, followed by the 4: 1 resonance, and so on. Therefore, this line of argument suggests that if the larva favours the transfer of energy between axial and transverse degrees of freedom, the transverse frequency should be at a low-order, even subharmonic of the axial frequency. For instance, given an axial characteristic frequency of *≈* 1 Hz, the strongest parametric driving of the transverse degrees of freedom should occur when the transverse frequency is 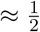, as observed in the real larva.

## E Analytical eigendecomposition of D_2,*c*_

We can find both **Φ**_*a*_ and **Λ**_*a*_ analytically by noting that **D**_2_ is a circulant matrix. Indeed, the *i*’th eigenvector of an arbitrary circulant matrix is given by [79]

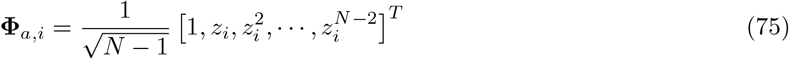

where we have used **Φ**_*a,i*_ to denote the *i*’th column of the eigenvector matrix **Φ**_*a*_, and 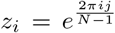 is the *i*’th element of the (*N −* 1)’th roots of unity, with 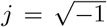 the imaginary unit. Using Euler’s complex exponential formula the *k*’th element of the *i*’th axial mode shape may be written

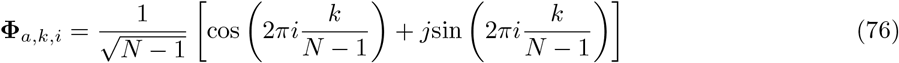

The real and complex parts of each vector can be considered as independent mode shapes, so that the modes thus come in pairs with identical spatial frequency,

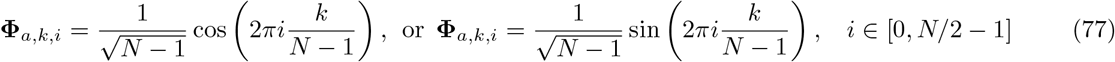

For an arbitrary (*N −* 1) × (*N −* 1) circulant matrix with entries

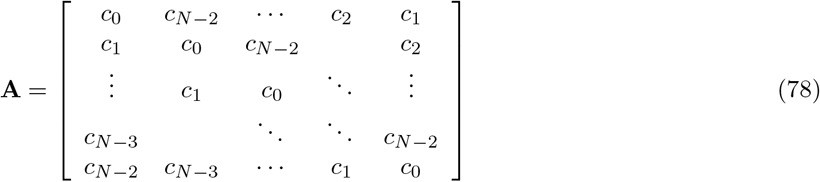

the eigenvalue corresponding to the *i*’th eigenvector is given by [79]

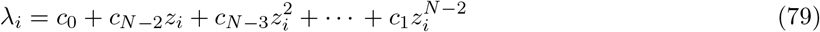

In the case of **D**_2_ we have *c*_0_ = 2 and *c*_1_ = *c*_*N−*2_ = *−*1, so that this reduces to

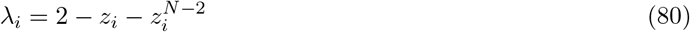

However, the *N −* 1’th roots of unity satisfy 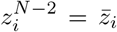, where the bar indicates the complex conjugate. Therefore,

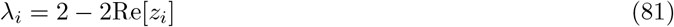

the real part of *z*_*i*_ can be found by using Euler’s complex exponential formula, yielding

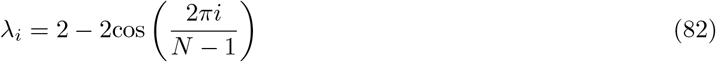

By using the trigonometric identity 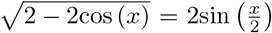 we may now calculate the natural frequency of the *i*’th axial mode

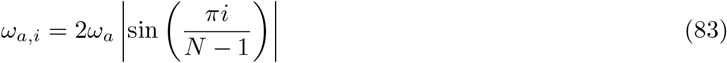

and the associated damping ratio

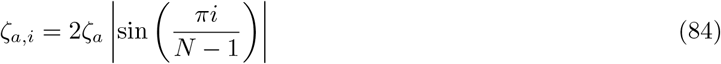

## F Analytical eigendecomposition of D_2,*f*_

Let us now turn our attention to the eigendecomposition of **D**_2,*f*_. The eigenvalue problem for this matrix can be stated as

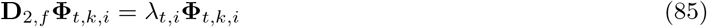

where the subscript *t* is intended to indicate that we are looking for the eigenvalue–eigenvector pairs related to *twisting* motions, *k* indexes over the segments of the model and *i* indexes over the eigenvector–eigenvalue pairs. For notational clarity we will drop the subscript *t* and index *i* until later in this subsection, and write the components of **Φ**_*t,k,i*_ as *v*_*k*_. We will define the index *k* to run from 0 at the tail to *n* + 1 at the head.

Using this notation, the eigenvalue problem may be written component-wise as

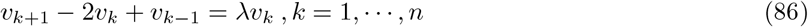

with the “boundary conditions”

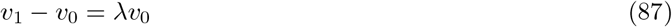

and

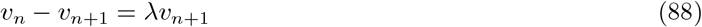

We note that for the boundary conditions to be satisfied in general we must have

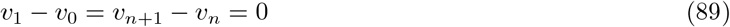

which motivates us to introduce new variables *w*_*k*_ = *v*_*k*+1_ *− v*_*k*_ in which the boundary conditions can be written simply as *w*_0_ = *w*_*n*_ = 0. We note that *v*_*k*+1_ *−* 2*v*_*k*_ + *v*_*k−*1_ = (*v*_*k*+1_ *−v*_*k*_) − (*v*_*k*_ *− v*_*k−*1_) so that the component-wise eigenvalue problem can be re-written

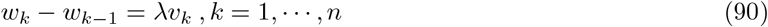

but *v*_*k*_ = *w*_*k−*1_ + *v*_*k−*1_, so

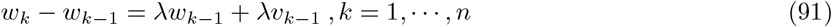

1228 Noting that *w*_*k−*1_ *− w*_*k−*2_ = *λv*_*k−*1_, we rewrite this as

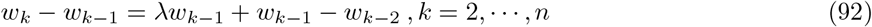

which can be rearranged to give

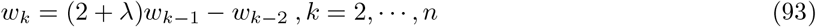

Introducing 2*α* = 2 + *λ* and shifting the index *k* by 1 gives the reccurrence relation

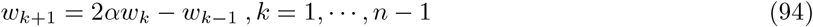

Considering *α* as an indeterminate, and assuming *w*_1_ = 1 (which we can always satisfy by normalising the eigenvector, provided that *w*_1_ ≠ 0), we may write

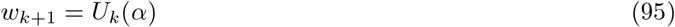

where *U*_*k*_ is the *k*’th Chebyshev polynomial of the second kind. Our boundary condition at the head then tells us

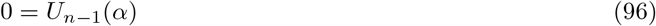

so that we can determine *α*, and thus the eigenvalues *λ*, by finding the roots of the (*n −* 1)’th Chebyshev polynomial of the second kind. These roots are well known, and tell us

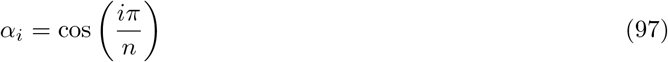

so that the *i*’th eigenvalue is determined by the equation

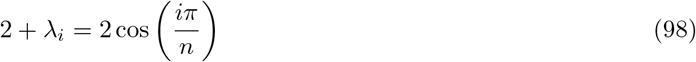

rearranging gives

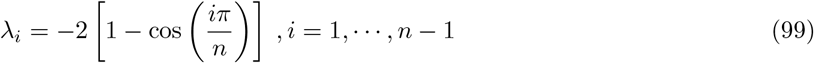

or, by a trigonometric identity,

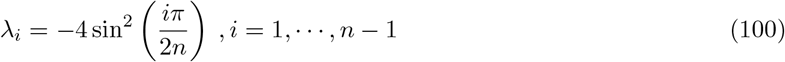

This gives us *n−*1 non-zero eigenvalues. There must clearly also be a zero eigenvalue corresponding to uniform rotation of all segments, with corresponding eigenvector given by *v*_*k*_ = *c* with *c* an arbitrary constant. We can incorporate the zero eigenvector into our expression above by writing

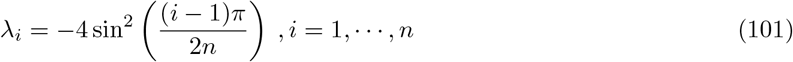

so that the torsional natural frequencies may be written

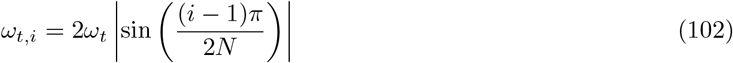

and the associated damping ratios are

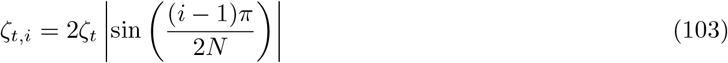

Each eigenvalue may now be substituted into the recurrence relations above in order to determine its corresponding eigenvector. Carrying out this procedure gives

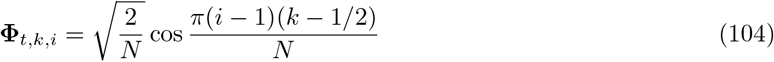

where the prefactor 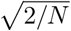 normalises the eigenvectors to unit length.

## G Analytical eigendecomposition of D_4_

To complete the eigendecomposition of our low-energy effective Hamiltonian, we now focus on the final eigenvalue problem

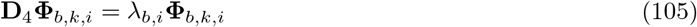

where the subscript *b* is intended to indicate that we are looking for the eigenvalue–eigenvector pairs related to *bending* motions, *k* indexes over the segments of the model and *i* indexes over the eigenvector–eigenvalue pairs. As in the previous section, for notational clarity we will drop the subscript *b* and index *i* until later in this subsection, and write the components of **Φ**_*b,k,i*_ as *v*_*k*_.

In the previous sections we exploited special properties of the second difference matrices to exactly compute their eigenvalues and eigenvectors. In particular, we exploited the circulant character of the second difference matrix with periodic boundary conditions to find its eigenvalues and eigenvectors using essentially a discrete Fourier transform, while we exploited the fact that the second difference matrix with free boundaries has a low bandwidth (non-zero elements only on the main diagonal and the two diagonals to either side, i.e. a bandwidth of 1) in order to write a tractable low-order recurrence relation for the eigendecomposition.

The fourth difference matrix we are now confronted with does not share these special properties – it is not circulant, and it has a higher bandwidth of 2 (i.e. five diagonals of the matrix have non-zero elements). This means that the discrete Fourier approach cannot be applied, and the recurrence relation will be of 5’th order and thus does not correspond to any Chebyshev polynomial.

Thus, we explore approximations to the eigendecomposition of **D**_4_. We begin by noting that our matrix can be decomposed into a product of rectangular *forward* and *backward* second difference matrices, i.e.

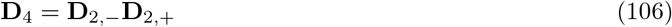

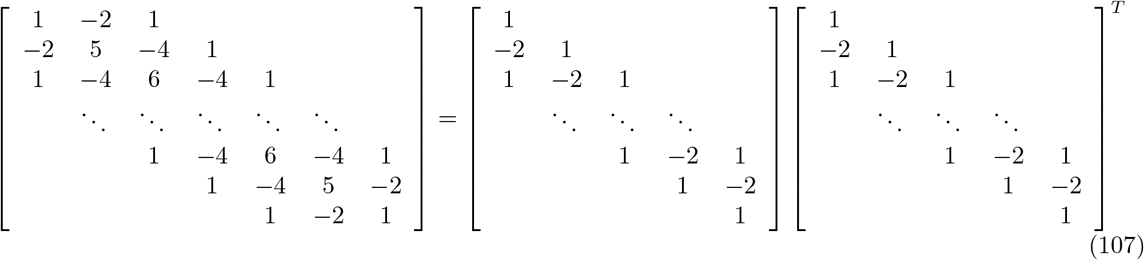

where we have noted that the backward and forward second difference matrices are simply related by taking the transpose, i.e. 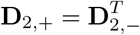. Since the nonzero eigenvalues of *AA*^*T*^ and *A*^*T*^ *A* are equal for any matrix *A*, this means that the nonzero eigenvalues of **D**_4_ = **D**_2,*−*_**D**_2,+_ must be equal to those of the product **D**_2,+_**D**_2,*−*_, i.e. the eigenvalues of

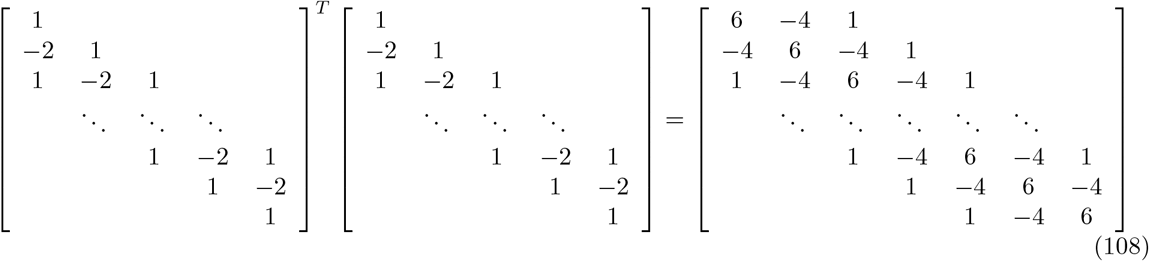

which is the (*N −* 2) × (*N −* 2) fourth difference matrix with fixed boundary conditions, which we denote **D**_4,fixed_. So far we have made no approximation, we have simply decomposed our original fourth difference matrix and proved that its nonzero eigenvalues must be identical to those for a related fourth difference matrix with different boundary conditions and a reduced dimensionality. Writing our problem in this way clarifies the mathematical issues we are facing – we need to find the eigenvalues of a real, symmetric, pentadiagonal Toeplitz matrix. There is currently no closed-form solution to this problem. However, we can use an asymptotics approach to find an exact solution in the limit *N* → *∞*.

To proceed, we approximate the (*N −* 2) × (*N −* 2) matrix **D**_4,fixed_ by its circulant counterpart, which we denote **D**_4,*c*_, with elements

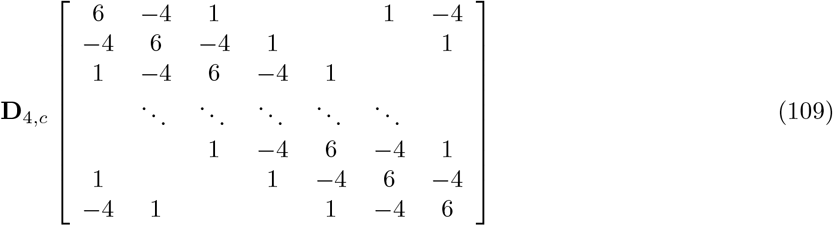

Clearly these two matrices are related by

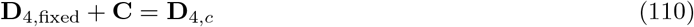

where the matrix **C** is zero everywhere apart from the six entries at the bottom left and top right corners (we choose the symbol **C** to stand for “corners”). Let us consider the relative error of this approximation as a function of the number of points *N* in our finite difference approximation. Clearly the element-wise approximation error is

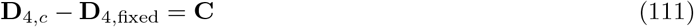

so that using a suitable matrix norm we can define a scalar measure of the absolute error

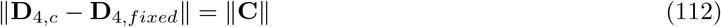

or the relative error

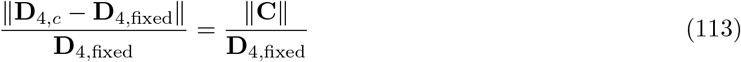

For concreteness let us use the *L*_2,1_ matrix norm, which is simply the sum of the Euclidean norms of the columns of a matrix. Since the matrix **C** only changes through inclusion of more zero elements as *N* grows, ∥ **C** ∥ is a constant in this norm. Meanwhile, incrementing *N* by 1 introduces an additional column into **D**_4,fixed_, always with the same 5 nonzero elements, so that the matrix norm should increase linearly as *N*→ *∞*. Therefore the relative error of our approximation should behave as *≈* 1*/N* as *N* increases, suggesting that the eigenvalues of the two fourth difference matrices should coincide aymptotically.

Next we note that the circulant fourth difference matrix can be factored into a product of circulant second difference matrices,

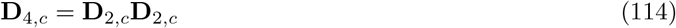

Performing eigendecomposition thus gives

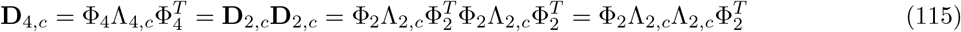

so that we may identify the eigenvalue matrix of the circulant fourth difference matrix with the product of eigenvalue matrices of the circulant second difference matrix Λ_4,*c*_ = Λ_2,*c*_Λ_2,*c*_. Since the eigenvalue matrices are by definition diagonal, this means that the *i*’th eigenvalue of the circulant fourth difference matrix is the square of the *i*’th eigenvalue of the circulant second difference matrix. Using our analytical solution for the *i*’th nonzero eigenvalue of the second difference matrix thus gives the approximation

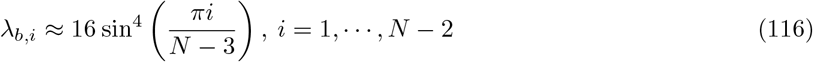

which is our final expression for the eigenvalues of the fourth difference matrix with free boundary conditions.

We can find the eigenvectors of **D**_4,*f*_ analytically. Let us start from the diagonalisation condition

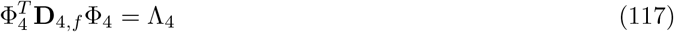

we then perform a change of coordinates according to Φ = **D**_1_Φ_4_ with **D**_1_ the backward first difference matrix. We invert the change of coordinates to give 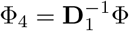, and insert into the diagonalisation equation to find

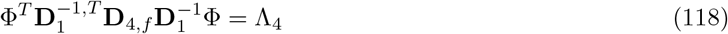

Thus we seek vectors Φ to diagonalise the central matrix product 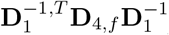. Once found, we will be able to obtain expressions for the eigenvectors of the original matrix via 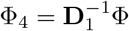.

We now note that our central matrix product gives us an augmented form of the (*N −* 1) × (*N −* 1) second difference matrix with free boundary conditions, i.e.

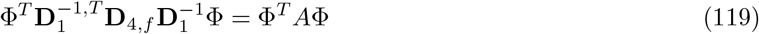

with the block diagonal matrix *A* given by

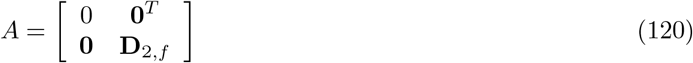

which must have eigenvectors [0, Φ_2_]^*T*^, where Φ_2_ is an (*N —* 1)-component eigenvector of the (*N —* 1) × (*N−* 1) second difference matrix with free boundaries. Thus, the eigenvectors of the fourth difference matrix with free boundaries are given by

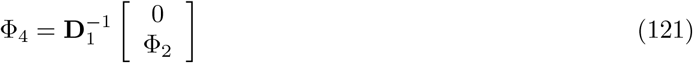

Meanwhile, the inverse of the backward first difference matrix is given by the *N* × *N* summing matrix

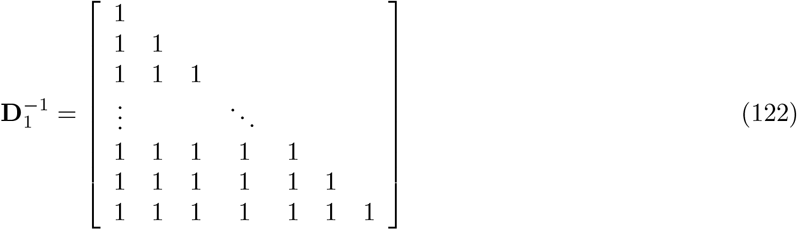

which can be used in conjunction with our analytical expressions for Φ_2_ from the previous section to find the analytical eigenvectors of **D**_4_. Carrying out this procedure gives the non-normalised eigenvectors

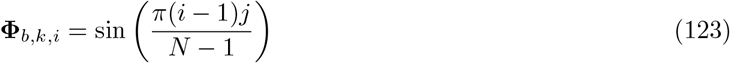

## H Detailed construction of the peristalsis/rolling model

It is marked that the axial and transverse modes come in pairs with identical frequencies. Each such pair of modes is called a *degenerate pair* due to their relationship to the degenerate eigenvalues of the difference matrices in the effective Hamiltonian. Note that the axial modes in a degenerate pair have identical spatial frequencies, while those in the transverse case have completely identical spatial components.

The degeneracy of the eigenvalues of the difference matrices in the effective Hamiltonian suggests that the choice of corresponding eigenvectors is somewhat arbitrary. Indeed, in principal we may choose any linear combination of the two degenerate eigenvectors to form the basis for each degenerate pair. Even if we limit ourselves to considering only orthonormal degenerate bases, we are still free to rotate the degenerate basis vectors through any angle *γ* that leaves them in the same plane. Such a transformation cannot change the mechanics, and so the *γ*-rotations of the degenerate bases constitute additional continuous symmetries of the effective theory. A beautiful result in classical physics known as Noether’s theorem tells us that these symmetries must be linked to additional conserved quantities. Let us make these ideas more concrete by considering a particular degenerate pair with modal coordinates *X*_1_, *X*_2_, conjugate momenta *p*_1_, *p*_2_, and natural frequency *ω*. To make the link between rotational symmetries and conservation laws most clear, we will briefly switch to using the Lagrangian framework of analytical mechanics. To do so, we begin with the Hamiltonian of the degenerate pair

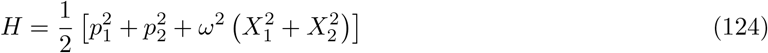

Taking the Legendre transform of the Hamiltonian yields the Lagrangian

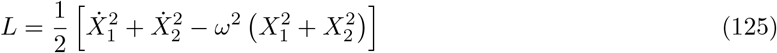

To make the rotational symmetry of this Lagrangian manifest, let us switch to polar coordinates *A, γ* defined via *X*_1_ = *A* cos *γ, X*_2_ = *A* sin *γ*. We will call *A* the degenerate amplitude and *γ* the degenerate phase. The Lagrangian is then

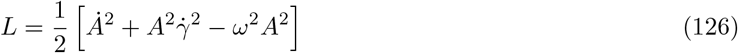

which clearly does not depend upon the angle *γ*. Indeed, the Euler-Lagrange equation for the degenerate phase gives

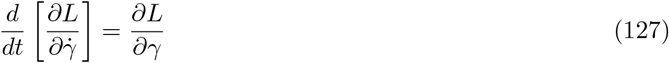

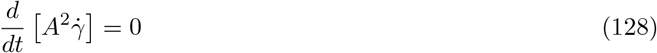

so that the bracketed expression on the left hand side must be conserved. Indeed, we recognise this expression immediately as a generalised form of angular momentum, which we term the degenerate angular momentum. We now switch back to using the Hamiltonian framework by taking the Legendre transform of the Lagrangian, finding the degenerate Hamiltonian in polar coordinates

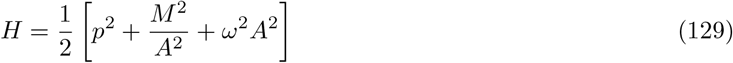

where 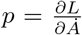 is the radial momentum conjugate to *A* and *M* is the degenerate angular momentum. This expression is simply a restatement of the total mechanical energy of the degenerate pair in polar coordinates. Note that we may now treat *M* as an arbitrary parameter and focus only on the dynamics of the degenerate amplitude *A* by introducing the effective potential energy 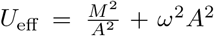. It should be clear that for a particular choice of degenerate angular momentum *M* there is now a minimum degenerate energy *E* = *H*(*p, A*) required for motion. Furthermore, the motion with this minimum energy corresponds to maintaining a constant degenerate amplitude, so that the trajectory of the system describes a circle in the original *X*_1_, *X*_2_ configuration space. For finite energies exceeding this minimum, the motion is instead confined to an annulus in the *X*_1_, *X*_2_ configuration space.

We now relate this picture back to the directly observable kinematics of the larva. First, we will convert from polar coordinates back to the degenerate modal coordinates *X*_1_, *X*_2_. To do so, we focus on the particular case in which *E* is set to its minimum value for a given *M*, corresponding to a conserved unit amplitude *A* = 1. We will set this value of *M* momentarily. Proceeding, we see that for our choice of *A*, 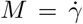 is simply the angular velocity of the motion, so that *γ* = *Mt* and the degenerate phase evolves linearly in time. Therefore, *X*_1_ = cos *Mt* and *X*_2_ = sin *Mt*. For this to coincide with our earlier results for the modal coordinates considered independently, we must have 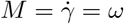, so that the modal coordinates *X*_1_ and *X*_2_ execute sinusoidal oscillations at the degenerate natural frequency *ω* with unit amplitude and a 90^*°*^ relative phase shift.

Next, we choose to interpret *X*_1_, *X*_2_ as the two modal coordinates of the *i*’th axial degenerate pair with natural frequency *ω*_*a,i*_. Using our expressions for the *i*’th pair of axial eigenvectors, we can write the axial displacement *x*_*k*_ of the *k*’th vertex of the midline as

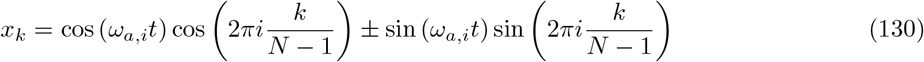

where we have dropped the normalising factor 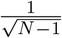. Using the identity cos (*a*) cos (*b*) ± sin (*a*) sin (*b*) = cos (*a* ± *b*), this further simplifies to

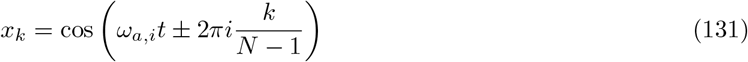

Interpreting 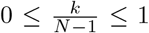 as a spatial coordinate ranging over the undeformed configuration of the body, this is in the form of a sinusoidal travelling wave, and the choice of a minus or plus sign in the argument corresponds to the choice of a forward- or backward-propagating wave, respectively.

Restricting our attention to the axial degenerate pair with lowest frequency and lowest dissipation, and further assuming that the segments of the larva should be held in place without slipping when not moving, the translational speed of the larva should be 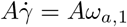.

Alternatively, we may choose to interpret *X*_1_, *X*_2_ as the two modal coordinates of a transverse degenerate pair. For instance, the pair of transverse modes with the lowest frequency, and lowest dissipation, corresponds to C-bending in the mediolateral and dorsoventral planes. The rotational symmetry of the degenerate basis in this case corresponds to our arbitrary choice of mediolateral/dorsoventral axes, and the corresponding degenerate angular momentum conservation law gives rise to the rotational propagation of a C-bend around the body of the larva with angular velocity *ω*_*b*,1_. Thus we see that there should be an exact correspondence between the angular velocity of rolling and the frequency of C-bending during unbiased behaviour, as we suggested in the main paper; this appears to be the case in the real animal.

In the presence of substrate constraints acting to hold the C-shaped bend in a fixed orientation in space (for instance the body may be held parallel to the plane of the substrate due to either “repulsive” ground-reaction forces or “attractive” surface tension forces), this motion would have to produce a compensatory rolling of the midline itself, at angular frequency *ω*_*b*,1_, in the opposite direction to the rotation of the C-shaped bend about the midline. In other words, in the presence of substrate interaction forces, we expect the conservation of the degenerate angular momentum to give rise to rolling behaviour. If we for now assume that the larva should roll without slipping, the translational speed of the larva would be 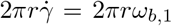 where *r* is the cross-sectional radius of the animal.

## I Modified Boltzmann distribution for degenerate modes

In this section we derive a maximum entropy distribution over the phase space of a pair of modes that share a degenerate (identical) frequency, subject to constraints of normalisation (probabilities must add up to 1), constant average energy, and constant generalised angular momentum *M*. To simplify the presentation we will work with sums over a discrete phase space and then take a continuum limit, rather than dealing explicitly with integrals [40]. The first part of the derivation consists simply of finding the classical Boltzmann distribution by maximising the Gibb’s entropy subject to the normalisation and energetic constraints. We then impose the constant generalised angular momentum constraint before calculating the partition function for the resulting modified phase space distribution.

The Gibb’s entropy that we will maximise is given by

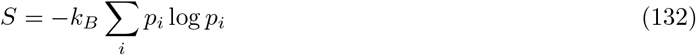

where the summation should be interpreted as ranging over *i* discrete states in the phase space. The maximisation is subject to the normalisation constraint

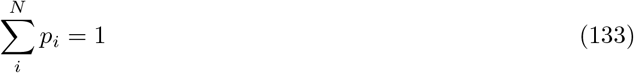

and the average energy constraint

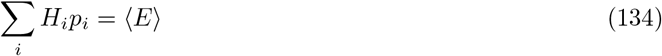

*H*_*i*_ is the Hamiltonian (total energy) calculated for the *i*’th state, which we constrain to take a constant average value ⟨*E⟩*. To find the constrained maximum of the entropy we use the method of Lagrange multipliers. We construct the objective function

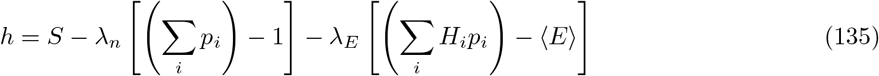

where we have introduced the Lagrange multipliers *λ*_*n*_ and *λ*_*E*_ associated with the normalisation and average energy constraints, respectively. To maximise *h* we set its partial derivative with respect to *p*_*i*_, *λ*_*n*_, and *λ*_*E*_ equal to zero, giving

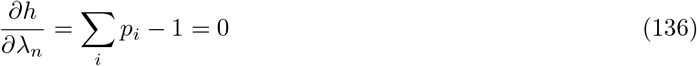

and

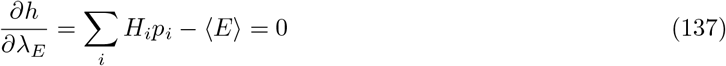

which simply recover our constraints, while taking the partial derivative with respect to the probabilities then gives

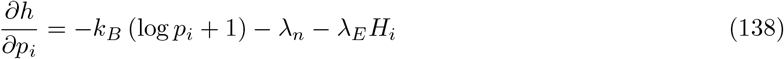

At a maximum of the Gibb’s entropy this quantity must equal zero. Setting this expression equal to zero and rearranging then gives the log-linear probability

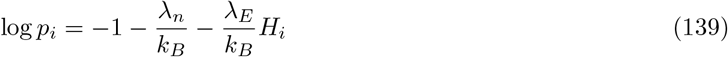

which we exponentiate to find the probability in factored form

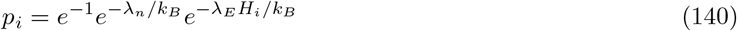

Clearly the first two factors are constant, and will be set by the normalisation condition via *λ*_*n*_. These factors thus correspond to the partition function, which we can write

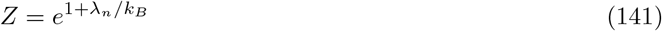

The remaining factor in the distribution will depend upon the average energy via the Lagrange multiplier *λ*_*E;*_, which we identify with the inverse temperature

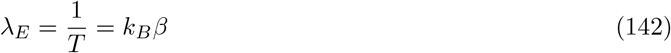

thus the phase space distribution in the absence of a conserved momentum *M* corresponds to the classical Boltzmann distribution for the canonical ensemble of classical statistical physics

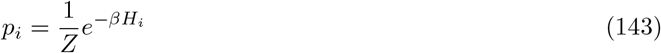

where the partition function *Z* can be determined via the normalisation constraint. In the continuum limit this becomes

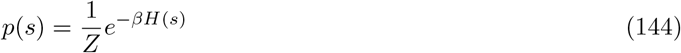

where *s* denotes a particular classical state in the phase space, consisting of canonical coordinates and their conjugate momenta. To take account of the exactly specified invariant momentum *M* we will make use of the degenerate Hamiltonian written in polar coordinates,

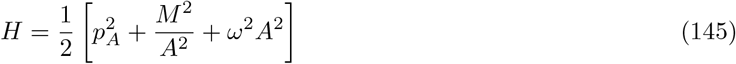

in which the degenerate angular momentum *M* usefully appears explicitly as a parameter, *A* is the degenerate amplitude and *p*_*A*_ is its canonically conjugate momentum. The degenerate phase space distribution is then

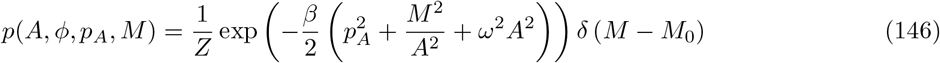

with *δ* the Dirac delta function. We can then write out our equation for the partition function *Z* by using the normalisation condition. We must replace the sum over discrete states in the normalisation condition with an integral over phase space, which ultimately gives us

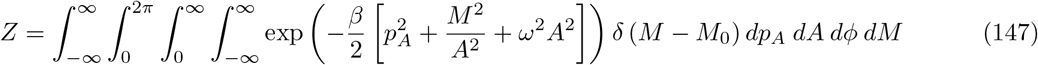

where the limits of integration are chosen to cover the phase space, i.e. *p*_*A*_ and *M* can take all values in the range *infty*, …, *∞, A* is constrained to be positive, and *ϕ* ranges from 0 to 2*π*.

Since the integrand is independent of *ϕ*, the integration over this variable simply contributes a factor 2*π*. Furthermore, due to the Dirac delta function enforcing our constraint *M* = *M*_0_, the integrand is zero for all *M* ≠ 0. Thus we can substitute the value *M* = *M*_0_ into the energy and integrate over the delta function, which simply contributes a factor of unity to the integral, so that we have

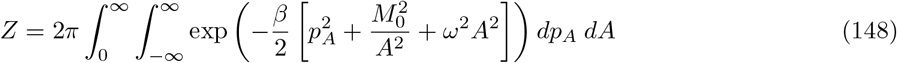

and we can focus our attention on the integration over *A* and *p*_*A*_. Since the kinetic energy (the first term in the integrand) depends only on *p*_*A*_ and does not depend upon *A*, and the effective potential (the second two terms in the integrand) depends upon *A* and does not depend upon *p*_*A*_, we can simplify further by factoring the integration as

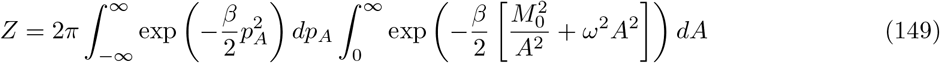

We now focus on the left-hand integral. The antiderivative can be written in terms of the error function, erf, as

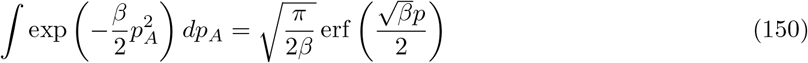

The error function tends to 1 as its argument tends to *∞* and tends to −1 as its argument tends to − *∞*, so the definite integral becomes

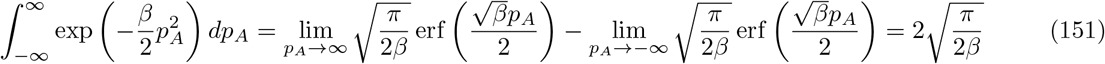

Substituting this into our expression for the partition function gives

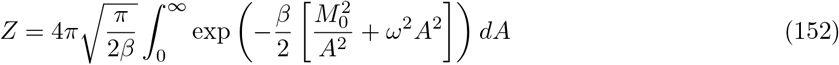

To calculate the remaining definite integral, we first calculate the antiderivative

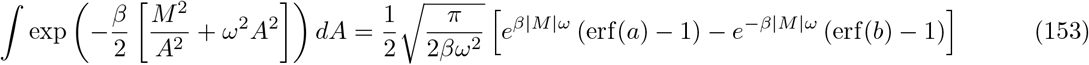

With

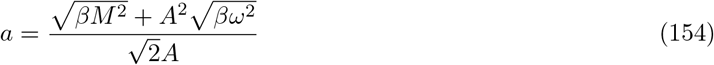

and

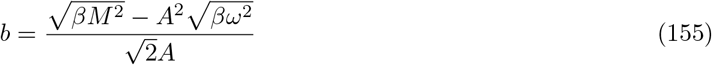

Clearly our integral must diverge at *A* = 0 since the integrand includes a term proportional to *A*^*−*^2, so to begin calculating our definite integral we instead find the limiting value of the antiderivative as *A* → 0 from above. We first distinguish two cases, corresponding to whether *M* is equal to zero or not. In the case *M* = 0 we have erf(*a*) → 0 and erf(*b*) → 0 as *A* → 0, so that the coefficients of the exponentials in the antiderivative are equal. However, the exponentials themselves are both equal to 1, since the exponents are 0, so that the exponentials cancel and the antiderivative must be equal to zero. Next considering the case *M* ≠ 0 we have erf(*a*) → 1 and erf(*b*) → 1 as *A* → 0, so that the coefficients of the exponentials are both equal to zero. Thus we have

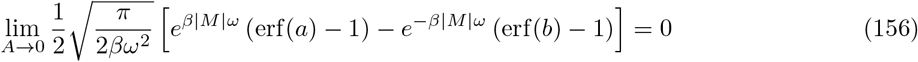

Now let us consider the upper limit in the definite integral, for which we must take the limit *A* → *∞* in the antiderivative. In this case erf(*a*) → 1 and erf(*b*) → *−*1 so that we have

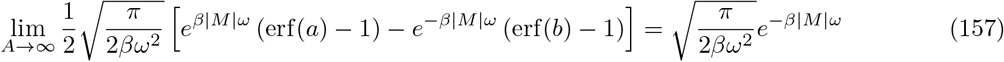

Given that the value of the antiderivative vanished at our lower limit of integration, this last expression at our upper limit must be equal to the final definite integral in the partition function. Substituting this expression into the partition function then gives

Substituting this into our expression for the partition function gives

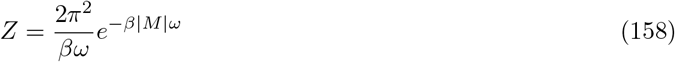

or, if we choose to measure phase space volumes in units of Planck’s constant *h*, as is customary in classical statistical mechanics [40], we have

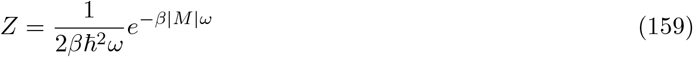

where *ħ* = *h/*2*π* is the reduced Planck’s constant. This is our final expression for the partition function of a degenerate pair of modes subject to an average energy constraint and the constraint of the conserved momentum *M* = *M*_0_.

Next, we compute the Helmholtz free energy from the partition function by using

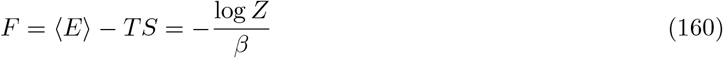

Which gives

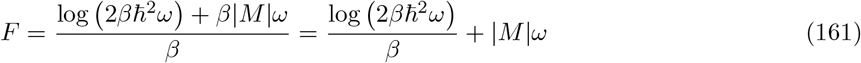

